# A TmaT-AftD interaction is required for the CmpL4 mycomembrane biogenesis pathway in *Corynebacterium glutamicum*

**DOI:** 10.64898/2026.07.31.741767

**Authors:** Anastacia R. Parks, Eric D. Snow, Victoria M. Marando, Michael J. James, Vincent de Bakker, Laura L. Kiessling, Suzanne Walker, Thomas G. Bernhardt

**Affiliations:** Department of Microbiology, Blavatnik Institute, Harvard Medical School, Boston, MA 02115, USA; Howard Hughes Medical Institute, Harvard Medical School, Boston, MA, 02115, USA; Department of Chemistry, Massachusetts Institute of Technology, Cambridge, MA 02139, USA

## Abstract

Bacteria in the Mycobacteriales order like mycobacteria and corynebacteria surround themselves with a multilayered cell envelope. Their cytoplasmic membrane is fortified by a peptidoglycan cell wall that is decorated with branched arabinogalactan (AG) polymers. The AG glycans are further modified with mycolic acids to form an outer membrane. Biogenesis of this mycomembrane requires the transport of mycolates from their site of synthesis in the cytoplasmic membrane to the cell surface. How mycolate transport is controlled and coordinated with AG synthesis has remained unclear. Mycolate transport is mediated by the essential RND-family transporter MmpL3 in mycobacteria and a pair of related, partially redundant transporters called CmpL1 and CmpL4 in corynebacteria. The acetyltransferase TmaT has also been implicated in mycolate transport in both types of bacteria. In corynebacteria, it is required for the production of acetylated mycolates, and this modification has been proposed to promote mycolate transport via both CmpL transporters in the model organism *Corynebacterium glutamicum* (*Cglu*). Here, we reinvestigated the function of TmaT in *Cglu* and found that it and several factors encoded in the *tmaT* locus are specifically required for mycolate transport via the CmpL4 transporter pathway. Notably, one of these additional genes encodes the arabinosyltransferase AftD involved in AG biogenesis. TmaT and AftD were found to interact, and our results indicate that this interaction is required for acetylated mycolate production, mycolate transport via the CmpL4 pathway, and normal arabinan synthesis. Thus, the TmaT-AftD interaction may serve as a regulatory link connecting mycolate transport with AG biogenesis.

**SIGNIFICANCE:** Mycobacteriales bacteria, including pathogens like *Mycobacterium tuberculosis* (*Mtb*), have a complex cell surface comprising an inner membrane, a cell wall modified with arabinogalactan (AG), and an outer mycomembrane made of mycolic acids linked to AG polymers. Because these surface biogenesis pathways are targeted by frontline anti-*Mtb* drugs, there is great interest in elucidating their underlying mechanisms. Here, we identify an interaction between factors involved in mycomembrane (TmaT) and AG biosynthesis (AftD) in the model organism *Corynebacterium glutamicum.* We show that this interaction is important for proper surface biogenesis and may therefore function to coordinate mycomembrane assembly with AG synthesis. This and other potential regulatory connections controlling envelope biogenesis represent attractive targets for future antibiotic development.

## INTRODUCTION

The Mycobacteriales order of bacteria includes deadly pathogens like *Mycobacterium tuberculosis* (*Mtb*) and *Corynebacterium diphtheriae* as well as the model organisms *Mycobacterium smegmatis* (*Msmeg*) and *Corynebacterium glutamicum* (*Cglu*). These bacteria are surrounded by a complex, diderm cell envelope (**Fig. 1**) that serves as a formidable barrier against entry of antibiotics and other external insults^1,2^ Additionally, the biogenesis of this envelope is the target of several front-line therapies for *Mtb* infections such as ethambutol and isoniazid^3^. Thus, a better understanding of the mechanisms promoting cell surface assembly in the Mycobacteriales has the potential to reveal new vulnerabilities that can be targeted for the treatment of infections caused by *Mtb* and related organisms.

**Figure 1.**
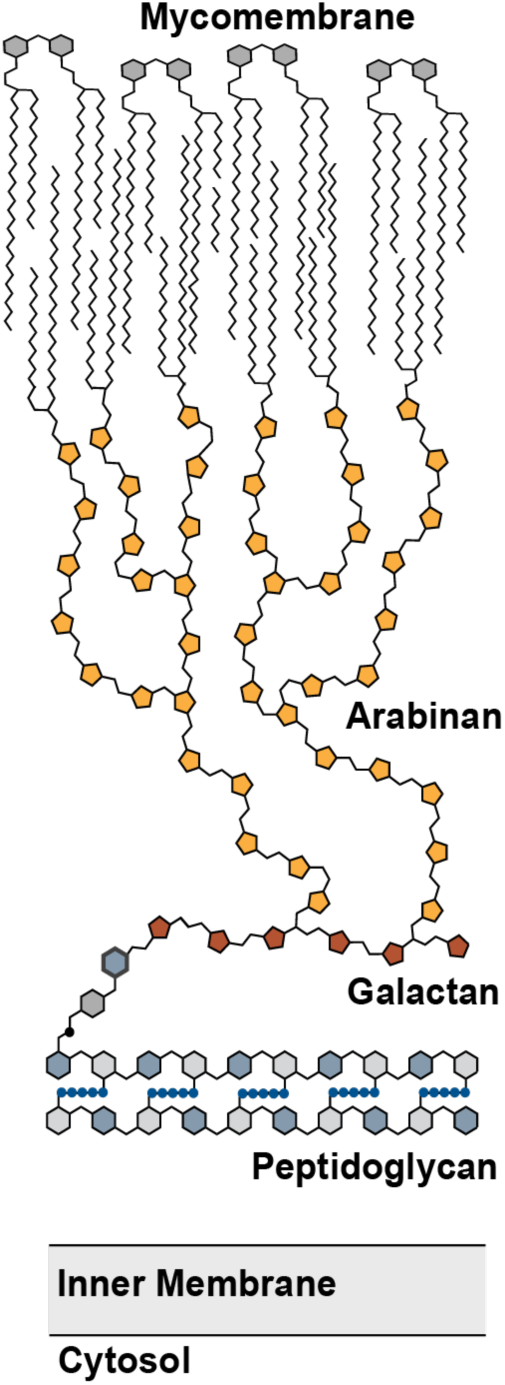
The cell envelope of Mycobacteriales bacteria. Shown is a schematic diagram of the cell envelope of corynebacteria and mycobacteria, highlighting the different layers: the inner (cytoplasmic) membrane, the peptidoglycan cell wall, the arabinogalactan layer, and the mycomembrane. For simplicity, lipoglycans like lipomannan and lipoarabinomannan are not shown. Figure was adapted from a previously published diagram^70^

Like most bacteria, the foundational layer of the mycobacterial/corynebacterial envelope is a peptidoglycan (PG) cell wall matrix that surrounds and fortifies the cytoplasmic membrane (**Fig. 1**). This matrix is further modified by the attachment of arabinogalactan (AG) polymers, which consist of a galactan backbone decorated with branched arabinan chains. The second membrane is composed of lipids called mycolic acids and is therefore referred to as the mycomembrane (**Fig. 1**)^2^. The inner leaflet of the mycomembrane is formed by mycolates covalently attached to the AG, and the outer leaflet is composed of the trehalose-linked mycolates trehalose monomycolate (TMM) and trehalose dimycolate (TDM). In corynebacteria, these molecules are often called trehalose corynemycolates (TMCM and TDCM), but we will refer to them as mycolates (TMM and TDM) here for simplicity^2,4^. ^5^Mycolic acids and the mycomembrane are essential for mycobacterial growth but are dispensable in *Cglu*. Thus, although cells lacking the mycomembrane have a severe growth defect, *Cglu* has proven to be a useful model system for studying the biogenesis of this unique membrane^6,7^.

The synthesis of mycolic acids begins in the cytoplasm and culminates with the production of TMM in the inner leaflet of the cytoplasmic membrane^6^. The TMM molecules are then thought to be flipped to the exterior face of the membrane by resistance, nodulation, and division (RND) transporters^8^. In mycobacteria, transport is mediated by the essential MmpL3 transporter whereas in corynebacteria the related CmpL1 and CmpL4 transporters play partially redundant roles in TMM transport^9,10^. Downstream steps of mycomembrane assembly likely require the extraction of TMM from the membrane. The mechanism of this extraction is not known but it also may involve the RND transporters^11^. Further transport is then thought to be carried out by mycoloyltransferases, which mobilize mycolate units to produce AG-linked mycolates, TDM in the mycomembrane, and mycoloylated proteins^12,13^. How the TMM supply is allocated to these distinct products and how mycomembrane biogenesis is generally coordinated with the synthesis of other envelope layers has remained poorly understood.

One way in which TMM utilization might be regulated is via its modification. Several years ago, the acetyltransferase TmaT was found to be required for the production of acetyl-TMM (^Ac^TMM) in *Cglu* (**Fig. 2A**)^4^. Mutants lacking TmaT accumulated TMM and were shown to be defective for TDM production, indicating a likely defect in mycolic acid transport. Accordingly, genetic profiling experiments indicate that TmaT is essential in *Msmeg* and *Mtb*^14–18^. This essentiality has since been confirmed in *Msmeg* where TmaT has been shown to be required for the biogenesis of TDM and AG-linked mycolates^19^. Thus, TmaT plays a critical and conserved role in mycomembrane biogenesis.

**Figure 2.**
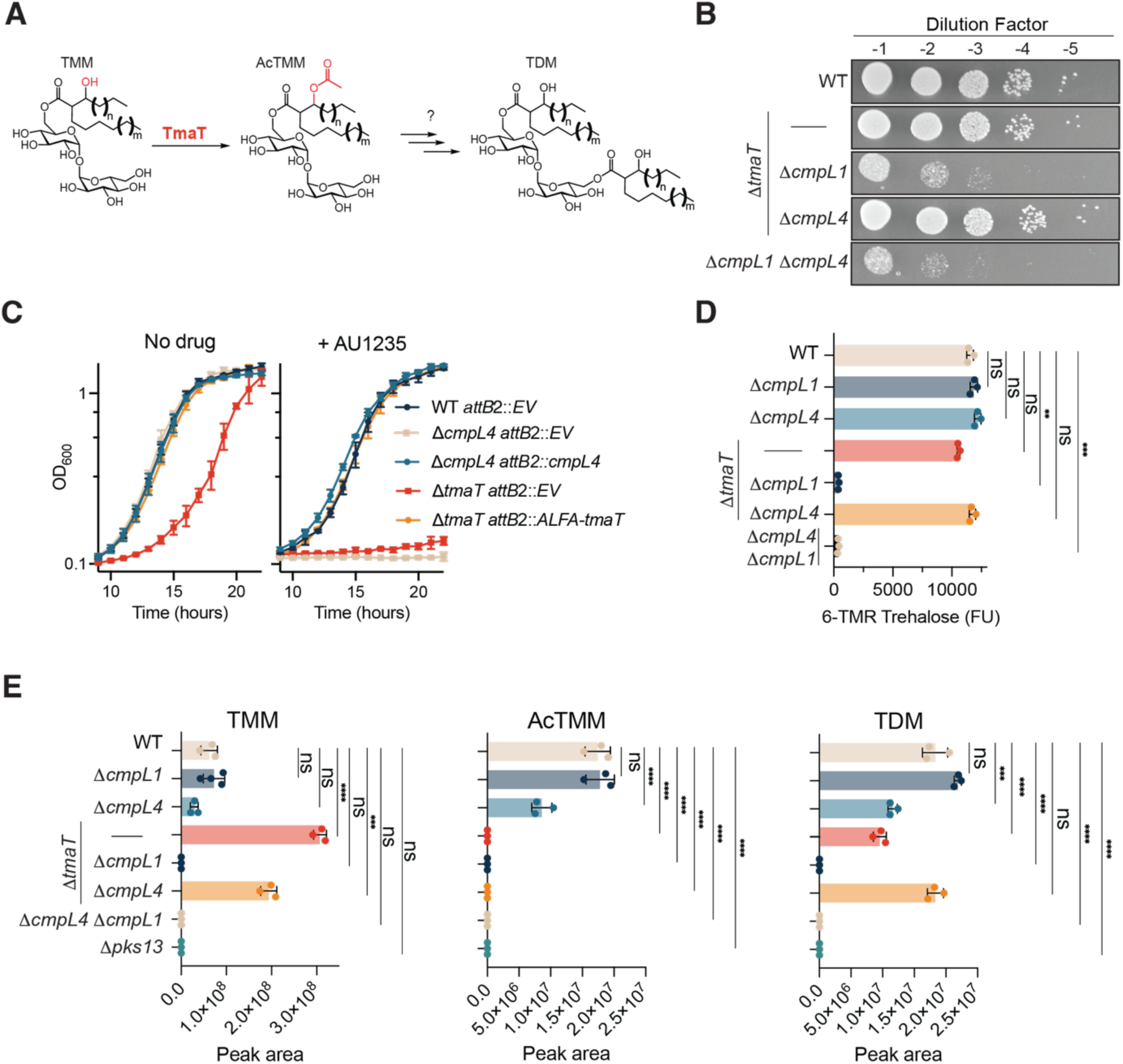
TmaT functions in the CmpL4-dependent mycomembrane biogenesis pathway. **A)** Schematic showing the TmaT-dependent modification of TMM to form ^Ac^TMM. Several steps in mycomembrane biogenesis are likely to occur after this modification, including potential deacetylation and formation of TDM. **B)** Cultures of strains MB001 [WT], H2654 [Δ*tmaT*], H4204 [Δ*cmpL1* Δ*tmaT*], H4203 [Δ*cmpL4* Δ*tmaT*], and H2708 [Δ*cmpL1* Δ*cmpL4*] were serially diluted, plated on BHIS (Brain-heart infusion plus sorbitol) and grown for 24 hours at 30°C before imaging. **C)** Cultures of strains H4896 [MB001 *attB2*::pARP116 (EV)], H5689 [Δ*cmpL4 attB2::pARP116 (EV)],* H5690 [Δ*cmpL4 attB2::cmpL4*], H5173 [Δ*tmaT attB2::pARP116 (EV)*], and H5424 [Δ*tmaT attB2::ALFA-tmaT*] were grown at 30°C in BHI in the presence or absence of AU1235 (25μM) in a 96-well microtiter plate. Growth was monitored by following the optical density at 600 nm (OD_600_). The experiment was performed in biological and technical triplicates. Error bars represent standard deviation of technical triplicates. Experiments were performed three independent times, and a representative result is shown. **D)** The strains MB001 [WT], H2654 [Δ*tmaT*], H4204 [Δ*cmpL1* Δ*tmaT*], H1476 [Δ*cmpL4*], H2692 [Δ*cmpL1*], H4203 [Δ*cmpL4* Δ*tmaT*], and H2708 [Δ*cmpL1* Δ*cmpL4*] were stained with 100 μM 6-TMR-Trehalose for 1 hour at room temperature. The OD_600_ and 6-TMR-Trehalose fluorescence were measured, and fluorescence units (FU) were calculated by dividing 6-TMR-Trehalose fluorescence by the OD_600_. Statistical comparisons were made using one-way ANOVA followed by Dunnett’s T3 multiple comparisons test with each mutant compared to WT (*** p<0.001, ** p<0.01, ns = not significant). Error bars represent standard deviation of technical replicates. **E)** LC-MS quantification of extracted trehalose monomycolate (TMM, C32:0), acetylated trehalose monomycolate (AcTMM, C32:0), and trehalose dimycolate (TDM, 2 x C32:0) from the indicated strains from (D) as well as H2261 [Δ*pks13*], which is defective for TMM synthesis. Lipid abundance was determined from the integrated peak area (ion counts) of the corresponding lipid species. Individual data points represent biological replicates (n=3), bars indicate the mean, and error bars represent the standard deviation. The experiment was performed three independent times. Statistical comparisons were made using one-way ANOVA followed by Dunnett’s T3 multiple comparisons test with each mutant compared to WT (*** p<0.001, **** p<0.0001, ns= not significant).

Mycolate transport by both the CmpL1 and CmpL4 in *Cglu* has been proposed to require acetylation of TMM by TmaT^4^. However, it has also been suggested that only one of the two transporters might require ^Ac^TMM as a substrate and that this specificity might account for the severity difference in phenotypes caused by TmaT inactivation in *Msmeg* versus *Cglu*^19^. We therefore sought to clarify the role of TmaT in mycolic acid transport in *Cglu*. Our genetic results indicate that TmaT is specifically required for the CmpL4 mycomembrane biogenesis pathway. Additionally, we show that several of the conserved genes in the *tmaT* locus previously implicated in mycolic acid transport also specifically function in this pathway^20,21^. We further discovered that the *aftD* gene, which encodes an arabinosyltransferase involved in AG biogenesis, is also required for a functioning CmpL4 pathway^22,23^. Finally, TmaT and AftD were found to form a complex, and this interaction was shown to be required for ^Ac^TMM production, proper mycomembrane assembly, and normal arabinan synthesis. Because this complex connects an enzyme implicated in TMM modification with an arabinosyltransferase, we propose that it forms part of a mechanism linking AG synthesis and mycomembrane biogenesis to ensure uniform surface growth. A contemporaneous, independent study of TmaT and AftD in *Msmeg* reached similar conclusions, indicating that this interaction and its role in envelope biogenesis is conserved^24^.

## RESULTS

### TmaT is required for the CmpL4 mycomembrane biogenesis pathway

Mutants lacking CmpL1 or CmpL4 individually do not display strong growth or mycomembrane biogenesis defects^25^. However, the simultaneous inactivation of both transporters is known to completely block TMM synthesis and mycomembrane assembly, resulting in a severe growth phenotype^25^. Therefore, to determine if TmaT functions in a specific pathway for mycomembrane biogenesis, we assessed the growth phenotype caused by *tmaT* deletion in combination with a deletion of either *cmpL1* or *cmpL4*. Cells inactivated for TmaT alone grew normally on solid medium and displayed a minor growth defect in broth culture (**Fig. 2B and 2C**). Growth did not appear to be strongly affected in cells lacking TmaT and CmpL4 (**Fig. 2B**). However, the simultaneous deletion of *tmaT* and *cmpL1* resulted in a severe growth defect that resembled that of cells lacking both CmpL1 and CmpL4 (**Fig. 2B**). Additionally, cells lacking TmaT showed a sensitivity to the CmpL1/MmpL3 inhibitor AU1235 that mirrored that of a similarly treated Δ*cmpL4* mutant (**Fig. 2C**)^9,10,26^. To monitor the status of mycolic acid synthesis and transport in the Δ*tmaT* Δ*cmpL1* mutant, we used the fluorescent trehalose analog 6-TAMRA-Trehalose (6-TMR-Tre), which is mycoloylated by mycoloyltransferases, forming labeled glycolipid in the mycomembrane^27^. Mutants individually inactivated for TmaT, CmpL1, or CmpL4 displayed wild-type levels of 6-TMR-Tre incorporation. By contrast and as expected, 6-TMR-Tre incorporation was undetectable in cells lacking both CmpL1 and CmpL4 (**Fig 2D**). A similar lack of 6-TMR-Tre labeling was observed for cells deleted for *tmaT* and *cmpL1* whereas cells deleted for *tmaT* and *cmpL4* labeled normally (**Fig 2D**).

To complement the 6-TMR-Tre labeling results, we measured the relative levels of TMM, ^Ac^TMM, and TDM in chloroform:methanol total lipid extractions from the single and double mutants using liquid chromatography and mass spectrometry (LC-MS) analysis. In agreement with previous studies, TmaT inactivation resulted in an increase in TMM accumulation, a complete loss of ^Ac^TMM species, and a decrease in TDM (**Fig. 2E** and S1-S3)^4,28^. A similar profile of TMM and TDM species was observed for the Δ*tmaT* Δ*cmpL4* mutant. However, like the Δ*cmpL1* Δ*cmpL4* mutant, TMM was undetectable in the Δ*tmaT* Δ*cmpL1* mutant (**Fig. 2E** and S1-S3, Table S1). Based on these results, we conclude that TmaT is specifically required for the proper functioning of the CmpL4 pathway for mycomembrane biogenesis.

### Genes in tmaT locus also participate in the CmpL4 mycomembrane biogenesis pathway

TmaT is encoded in a conserved genetic locus that includes two additional genes, *mtrP* (*cgp_3168*) and *mmpA* (*cgp_3165*), that have been previously implicated in mycomembrane biogenesis^29,30^ (**Fig. 3A**). MtrP is a putative methyltransferase and MmpA is an integral membrane protein of poorly understood function. Like cells inactivated for *tmaT*, mutants deleted for *mtrP* and *mmpA* were found to have defects in ^Ac^TMM production^29,30^. They were also shown to accumulate TMM and to be defective for TDM production, suggesting that TmaT activity and mycolic acid transport were impaired by their inactivation. Accordingly, both genes are essential for mycobacterial growth^31–35^. Notably, *mtrP* and *mmpA* along with the nearby genes of unknown function *cgp_3166* and *cgp_3167* but not *cgp_3164* (*lmcA*) were found to have phenotypic profiles that are highly correlated with *tmaT* (**Fig. 3B**)^36^. We also observed a correlation with *cmpL4* albeit not as strong as the other loci, possibly because the phenotypic profile of *cmpL4* for certain conditions has more summed read counts than *tmaT* **(Fig. S4**). Nevertheless, we wondered whether these additional factors encoded in the *tmaT* locus are also required for the CmpL4 mycomembrane biogenesis pathway. Individual deletions of *mmpA*, *mtrP*, *cgp_3166*, and *cgp_3167* were combined with a deletion of either *cmpL1* or *cmpL4* and the growth phenotype of the resulting mutants was assessed. All the double mutants lacking *cmpL4* grew normally (**Fig. 4A**). However, most of the combinations with a *cmpL1* deletion displayed a severe growth defect that resembled that of Δ*cmpL1* Δ*cmpL4* and Δ*cmpL1* Δ*tmaT* double mutants. The exception was the Δ*cmpL1* Δ*cgp_3167* double mutant, which grew like wild type (**Fig. 4A**). Additionally, 6-TMR-Tre incorporation was blocked in all double mutants lacking CmpL1 except for the Δ*cmpL1* Δ*cgp_3167* double mutant, which labeled similarly to wild-type **(Fig 4B).** Finally, all single mutants except for Δ*cgp_3167* displayed a slight growth defect when grown in liquid rich medium and a sensitivity to the CmpL1 inhibitor AU1235 **(Fig 4C).** We therefore conclude that all genes in the *tmaT* locus except for *cgp_3167* and *cgp_3164*, which have previously been shown to be involved in lipoglycan synthesis^37,38^, are required for the CmpL4 pathway for mycomembrane biogenesis.

**Figure 3.**
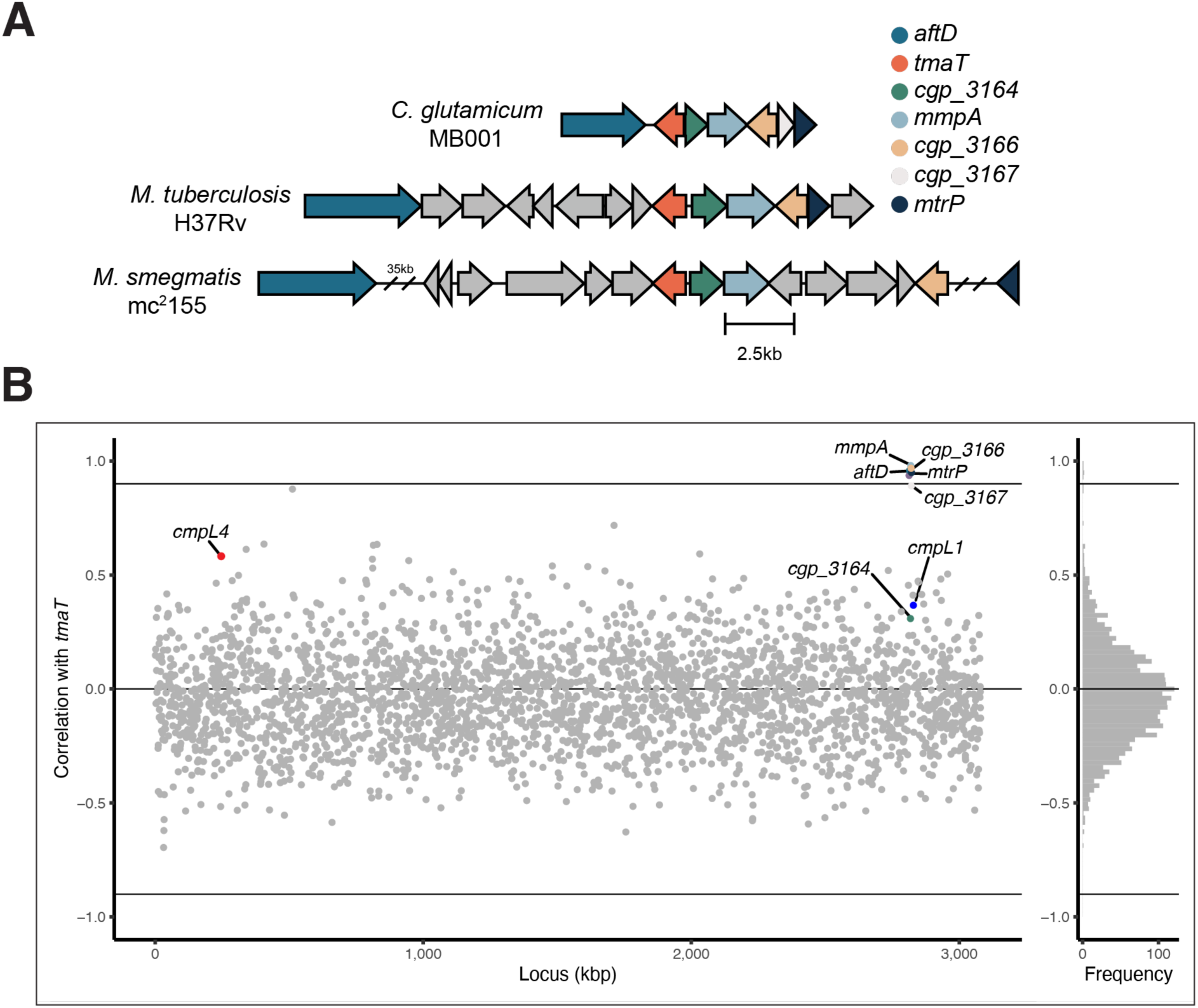
The *tmaT* locus and genes correlated with *tmaT* via phenotypic profiling. **A)** Genomic context of *tmaT* in *Cglu* strain MB001, *M. tuberculosis* strain H37Rv, and *M. smegmatis* strain mc²155. Arrows represent predicted ORFs, oriented according to the direction of transcription. Genes conserved across all three loci are colored as indicated in the legend: *aftD*, *tmaT*, *cgp_3164*, *mmpA*, *cgp_3166*, *cgp_3167*, and *mtrP*. Gray arrows denote flanking ORFs outside this conserved set. Loci are drawn to scale except for two regions in *M. smegmatis* (denoted “35 kb” and “//”), which are not to scale. Scale bar, 2.5 kb. **B)** Genome-wide correlation of phenotypic profiles with the profile of *tmaT*. Data is from a prior study^71^. Each point is a gene plotted by genomic locus position (kbp) and Pearson’s correlation coefficient with the phenotypic profile of *tmaT*. Horizontal line indicates |*r*| = 0.9, exceeded by the conserved locus genes from (**A**) — *aftD*, *mmpA*, *cgp_3166*, *cgp_3167*, and *mtrP* —indicating a shared phenotypic profile with *tmaT,* except for *cgp_3164*. *cmpL4*, which displays a moderately similar phenotypic profile to *tmaT* (see also Fig. S4), is shown as well. Right, histogram of the distribution of correlation coefficients across all genes screened.

**Figure 4.**
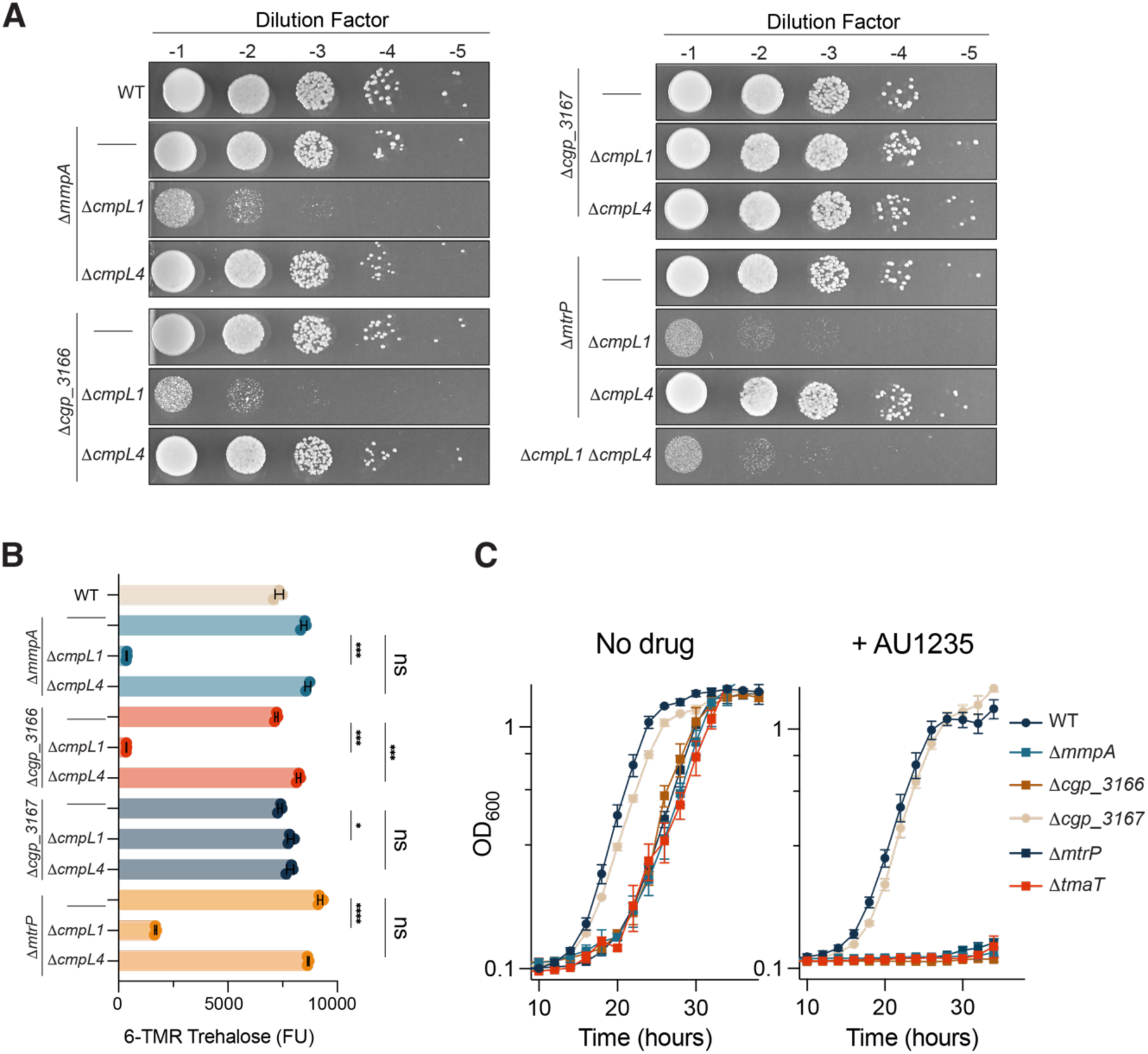
Factors encoded in the conserved *tmaT* locus are required for function of the CmpL4 pathway. **A)** Cultures of strains MB001 [WT], H2864 [Δ*mmpA*], H6042 [Δ*cmpL1 mmpA*::*kan*^+^], H6041 [Δ*cmpL4 mmpA*::*kan*^+^], H5713 [Δ*cgp_3166*], H6044 [Δ*cmpL1 cgp_3166*::*kan*^+^], H6043 [Δ*cmpL4 cgp_3166*::*kan*^+^], H6045 [*3167*::*kan*^+^], H6047 [Δ*cmpL1 cgp_3167*::*kan*^+^], H6046 [Δ*cmpL4 cgp_3166*::*kan*^+^], H4219 [Δ*mtrP*], H4347 [Δ*cmpL1 mtrP*::*kan*^+^], and H4346 [Δ*cmpL4 mtrP*::*kan*^+^] were serially diluted, plated on BHI and grown for 24 hours at 30°C before imaging. **B)** Strains from (**A**) were labeled with TMR-Tre and incorporation was determined as in **2D**. The results for each single deletion mutant were compared to the corresponding Δ*cmpL1* and Δ*cmpL4* double mutant by Welch’s ANOVA followed by pairwise comparisons of each single mutant to its corresponding Δ*cmpL1* or Δ*cmpL4* double mutant using Dunnett’s T3 multiple comparisons test (8 comparisons, α=0.05, *<0.05, *** p<0.001, **** p<0.0001, ns= not significant). Error bars represent standard deviation of biological and technical replicates. **C)** The indicated strains from (**A**) were grown in the presence or absence of AU1235 and growth monitored as in **2C**. Experiments were performed in technical and biological triplicate three independent times. Error bars represent standard deviation.

### The AG synthesis enzyme AftD plays a role in mycomembrane biogenesis

Another gene found to be highly correlated with *tmaT* by phenotypic profiling was *aftD* (*cgp_3161*), which is also located adjacent to *tmaT* but on the opposite side from the conserved genetic locus previously linked with TmaT function^29,30^ **(Fig. 3A).** AftD encodes an arabinofuranosyltransferase (AraT) involved in AG branching in concert with the arabinofuranosyltransferases AftA (adds single D-Ara*f* to C5 of D-Gal*f*), Emb (α-(1→5) extension), AftC (α-(1→3) branching), and AftB (β-(1→2) capping) (**Fig. 5A**).^39–45^ It is thought that AftD participates in late-stage polymerization of lipoarabinomannan (LAM) and AG branches as either an α-(1→3) and/or α-(1→5) linking enzyme, however its precise role in AG/LAM biogenesis has yet to be determined.^46,47^ As with most of the other genes in the *tmaT* locus, the combination of an *aftD* deletion with Δ*cmpL1* but not Δ*cmpL4* resulted in a severe growth defect resembling that of Δ*cmpL1* Δ*cmpL4* and Δ*cmpL1* Δ*tmaT* double mutants (**Fig. 5B**). No growth phenotype was observed when *tmaT* and *aftD* were simultaneously inactivated (**Fig. 5B**). The Δ*cmpL1* Δ*aftD* double mutant did not incorporate 6-TMR-Tre as observed with the Δ*cmpL1* Δ*cmpL4* mutant **(Fig. 5C).** Thus, like TmaT and the other factors encoded in the *tmaT* locus, AftD is required for the function of the CmpL4 mycomembrane biogenesis pathway. To determine whether this role for AftD is specific or the result of a general defect in AG branching, we compared the effect of an *aftD* deletion with that caused by the inactivation of AftC, the enzyme thought to be upstream of AftD in the AG biogenesis pathway that initiates the formation of arabinan branches (**Fig. 5A**)^48^. If any arabinan-branching defect impaired the CmpL4 pathway, cells lacking AftC should phenocopy those lacking AftD. As expected, mutants lacking AftD were sensitive to the CmpL1 inhibitor AU1235 (**Fig. 5D**). However, cells lacking AftC, which are also defective for AG branch formation, did not display a similar sensitivity (**Fig. 5E**). Thus, our results suggest that AftD plays a role in mycomembrane biogenesis that is distinct from its typical function in AG synthesis.

**Figure 5.**
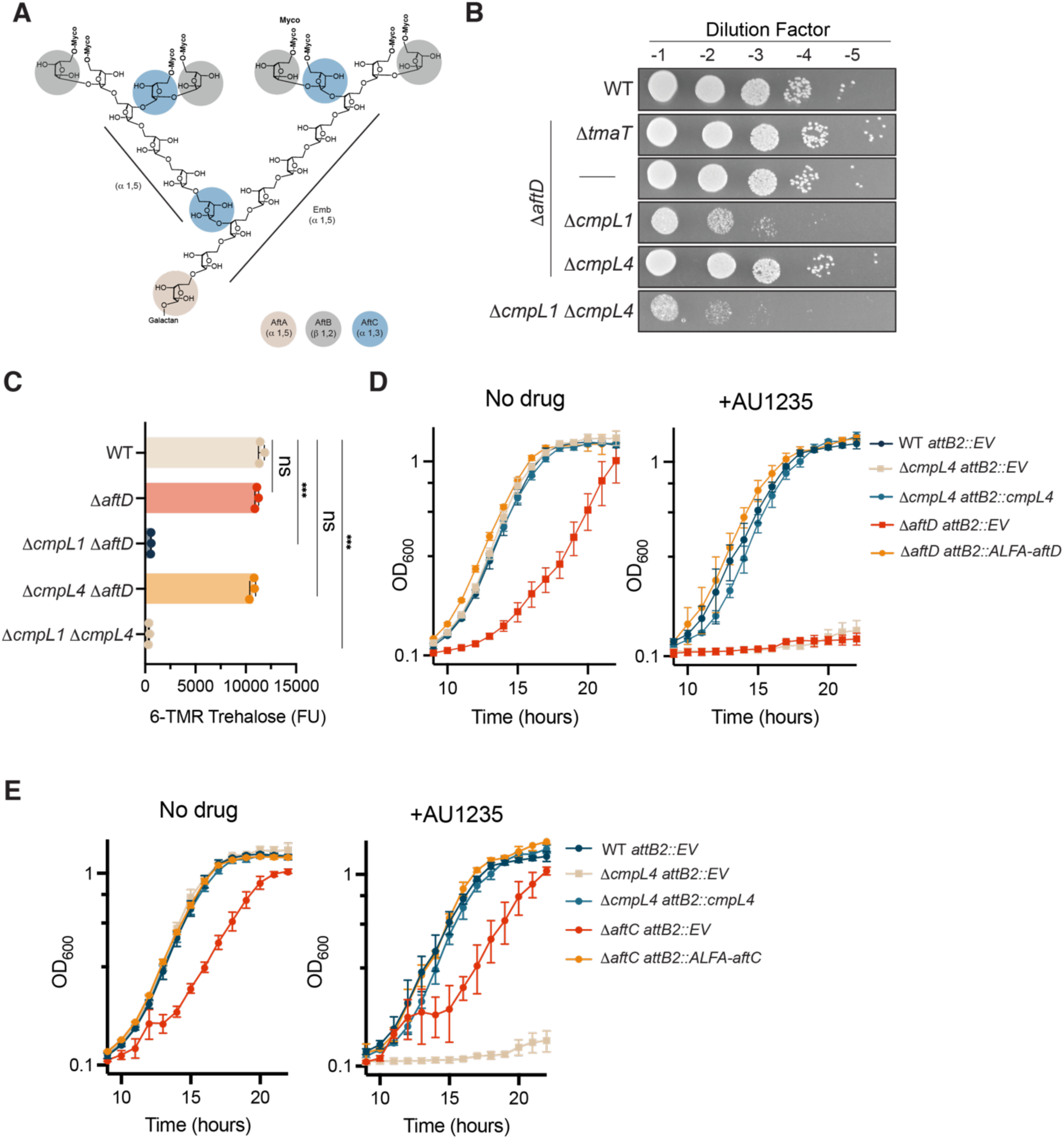
AftD is also required for the CmpL4 transport pathway. **A)** Schematic of the arabinogalactan (AG) arabinan domain, depicting the branching architecture and the sites of action of arabinosyltransferases AftA, AftB, AftC, and Emb (EmbA/B/C), with the designated direction of addition of arabinosyl residues added by each enzyme indicated in parentheses. Activities shown were based on previous work^39–45^. **B)** Cultures of strains MB001 [WT], H2657 [Δ*aftD*], H2657 [Δ*tmaT*], H4857 [Δ*tmaT* Δ*aftD*], H5520 [Δ*cmpL1* Δ*aftD*], H5436 [Δ*cmpL4* Δ*aftD*], and H2708 [Δ*cmpL1* Δ*cmpL4*] were serially diluted, plated on BHI and grown for 24 hours at 30°C before imaging. **C)** Strains from (**B**), except H2657 and H4857, were labeled with TMR-Tre and incorporation was determined as in **2D**. Statistical comparisons were made using Welch’s ANOVA followed by pairwise comparisons of each mutant to WT using Dunnett’s T3 multiple comparisons test (4 comparisons, α=0.05, *** p<0.001, ns = not significant). Data represent biological replicates with mean shown. Experiment was conducted three separate times. **D)** Cultures of strains H4896 [MB001 *attB2*::pARP116 (EV)], H5689 [Δ*cmpL4 attB2::pARP116* (EV)], H5690 [Δ*cmpL4 attB2::cmpL4*], H4873 [Δ*aftD attB2::pARP116* (EV)], and H4861 [Δ*aftD attB2::ALFA-aftD*] were grown in the presence or absence of AU1235 and growth monitored as in **2C**. Experiments were performed in technical and biological triplicate three independent times. Error bars represent standard deviation. **E)** Cultures of strains H4896 [MB001 *attB2*::pARP116 (EV)], H5689 [Δ*cmpL4 attB2::pARP116* (EV)], H5690 [Δ*cmpL4 attB2::cmpL4*], H5457 [Δ*aftC attB1::pARP116* (EV)], and H5444 [Δ*aftC attB1::ALFA-aftC*] were grown in the presence or absence of AU1235 and growth monitored as in **2C.** Error bars represent standard deviation. Experiments were performed in technical and biological triplicate, and a representative example is shown.

### Cells lacking AftD have similar envelope changes to those inactivated for TmaT

It was previously shown that inactivation of *mtrP* or *mmpA* results in changes to mycolic acid synthesis that mirror those observed in Δ*tmaT* cells^29,30^. Given that phenotypic profiling also links *aftD* with *tmaT*, we investigated whether mutants lacking *aftD* also display changes in the trehalose mycolate pools similar to those observed in cells inactivated for TmaT. Total lipids were extracted from wild type, Δ*tmaT,* and Δ*aftD* cells and analyzed by LC/MS to detect TMM, ^Ac^TMM, and TDM (**Fig. 6A**). As observed previously, Δ*tmaT* cells accumulated TMM relative to wild type (**Fig. 6A**). They were also devoid of ^Ac^TMM and displayed reduced levels of TDM relative to wild type cells (**Fig. 6A**), suggestive of a mycolic acid transport defect. This altered lipid profile was restored to normal by the production of an ALFA-tagged TmaT variant from a chromosomally integrated expression construct (**Fig. 6A**). Cells lacking AftD displayed a similarly altered profile of trehalose mycolates, most notably the complete loss of ^Ac^TMM species. Importantly, these alterations were also corrected by the production of an ALFA-tagged variant of AftD from an ectopic locus (**Fig. 6A**). To assess the effect of TmaT or AftD inactivation on AG synthesis, we used the biosynthetic probe 2-AzFPA (2-Azido-(Z,Z)-farnesyl phosphoryl-β-D-arabinose)^49^. The modified D-arabinose in the probe is incorporated into the arabinan layer following its addition to live cells. Probe incorporation is then detected using click chemistry to add a fluorescent label followed by flow cytometry detection^49^. Notably, the Δ*tmaT* and Δ*aftD* mutants both exhibited an increase in 2-AzFPA labeling as compared to wild type and complemented strains, indicating that the mutations cause an alteration in the arabinan synthesis pathway (**Fig. 6B and Fig. S5**). Thus, the inactivation of TmaT or AftD results in correlated changes in the synthesis of both trehalose mycolates and arabinan polymers, suggesting that each protein may reciprocally influence the activity of the other.

**Figure 6.**
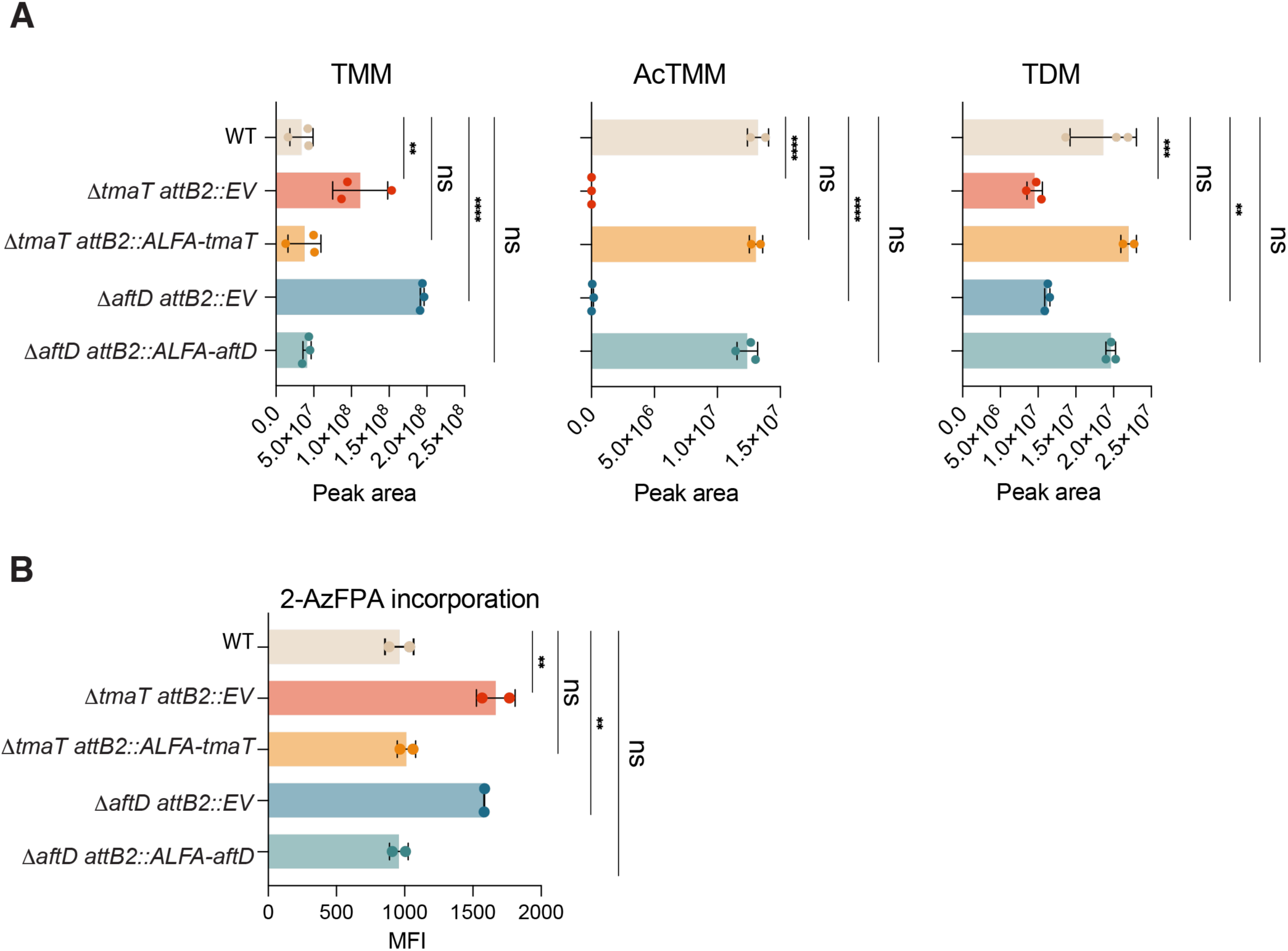
AftD inactivation phenocopies a TmaT defect. **A)** LC-MS quantification of peak areas for trehalose monomycolate (TMM, C32:0), acetylated trehalose monomycolate (^Ac^TMM, C32:0), and trehalose dimycolate (TDM, 2x C32:0) in strains H4896 [MB001 *attB2*::pARP116 (EV)], H5173 [Δ*tmaT attB2::pARP116 (EV)*], and H5424 [Δ*tmaT attB2::ALFA-tmaT*], H4873 [Δ*aftD attB2::pARP116 (EV)*], and H4861 [Δ*aftD attB2::ALFA-aftD*]. Statistical comparisons were made using one-way ANOVA with each mutant compared to WT (** p<0.01, **** p<0.0001, ns= not significant). Data represent individual biological replicates with mean shown. **B)** Flow cytometry analysis of 2-AzFPA (2-Azido-(Z,Z)-farnesyl phosphoryl-β-D-arabinose; 150μM) labeled strains from (**A**) treated with DBCO-AF647. Mean fluorescence intensity (MFI) was calculated using the geometric mean and one-way ANOVA with each mutant compared to WT (** p<0.01, ns= not significant). Data represent replicates with mean shown. Experiments were conducted at least two independent times.

### AftD and TmaT form a complex

The phenotypic similarities between mutants inactivated for TmaT and those lacking the other factors encoded within the *tmaT* locus suggested that one or more of these proteins may interact with TmaT to promote its activity. We therefore performed pairwise AlphaFold 3 predictions for complexes between TmaT and MmpA, Cgp_3166, Cgp_3167, MtrP, or AftD. The only high-confidence prediction was for an AftD-TmaT complex, with the two proteins predicted to interact in several other members of the Mycobacteriales (**Fig. 7A and S6-S7**)^50^. To validate this interaction, strains producing a Halo-tagged AftD and either untagged or an ALFA-tagged TmaT were constructed. Following anti-ALFA resin purification, Halo-AftD was specifically detected in the elution fractions from extracts producing ALFA-TmaT (**Fig. 7B**).

**Figure 7.**
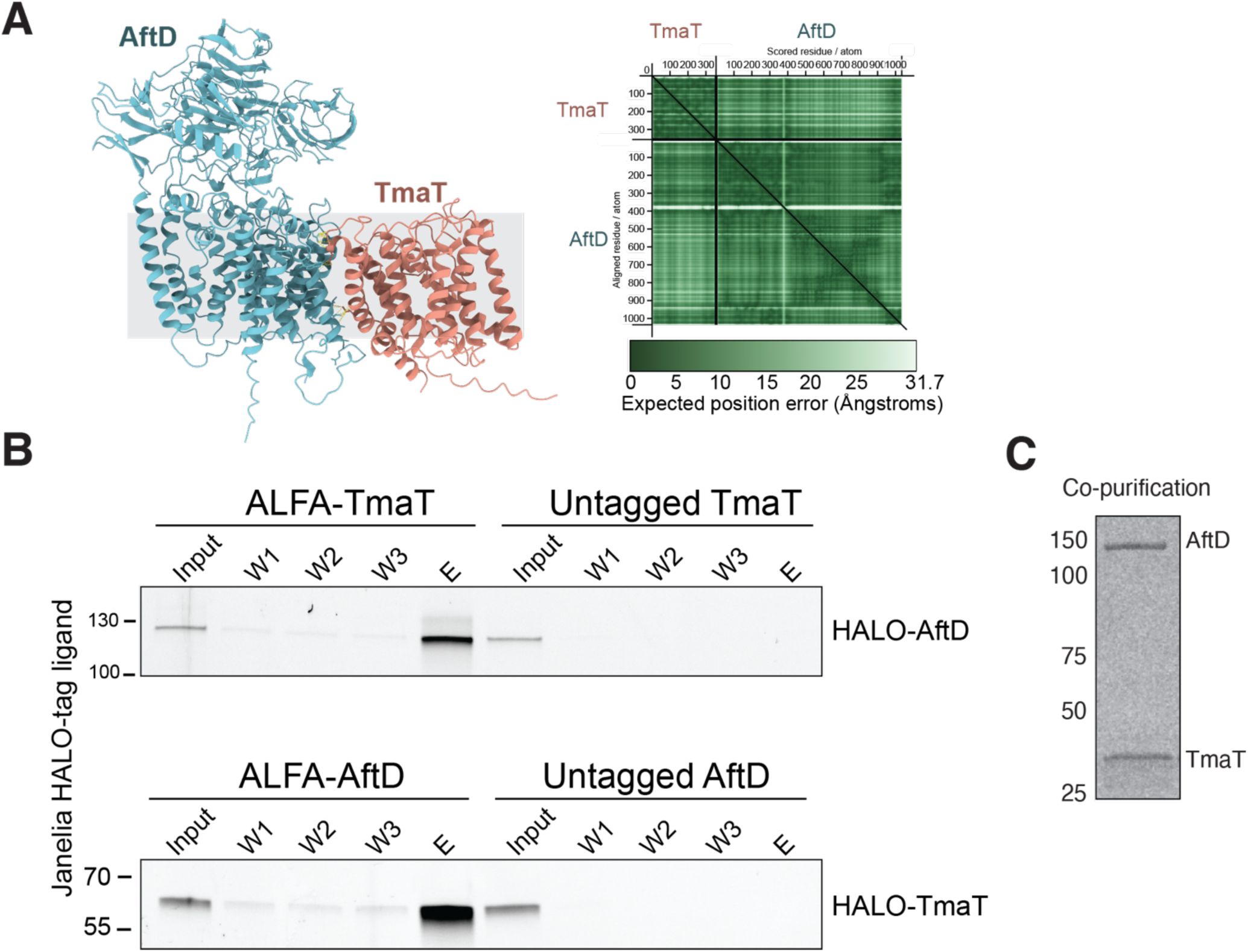
TmaT and AftD form a complex. **A)** Alphafold 3 predicted structure of the TmaT(salmon)-AftD(teal) complex (left) and corresponding PAE plot (right)^50,72^ with an iptM of 0.88. **B)** Co-immunoprecipitation results of HALO-AftD with ALFA-TmaT (top, strain H5426) and HALO-TmaT with ALFA-AftD (bottom, strain H5534) using ALFA-resin. Input, wash fractions (W1-W3), and elution (E) were treated with Janelia HALO-tag ligand (Fluor 549) and resolved by SDS-PAGE and imaged on a Typhoon imager using an Alexa 546 filter. Untagged TmaT [H5425] and untagged AftD [H5533] serve as negative controls. **C)** Coomassie-stained gel of co-purified FLAG-AftD and His-TmaT following co-expression in *E. coli*, showing co-elution of both proteins at their expected molecular weights.

Similarly, in strains producing Halo-TmaT and either untagged AftD or ALFA-AftD, Halo-TmaT was specifically detected in the anti-ALFA elution of cells producing ALFA-AftD (**Fig. 7B**). Additionally, we heterologously produced His_10_-TmaT and FLAG-tagged AftD in *Escherichia coli*, solubilized the membranes in detergent, and performed immobilized metal affinity chromatography followed by anti-FLAG affinity chromatography. This tandem purification procedure yielded what appears to be a 1:1 TmaT–AftD complex (**Fig. 7C**). We conclude that TmaT and AftD interact directly as predicted by AlphaFold 3.

### The TmaT-AftD interaction is required for the CmpL4 mycomembrane biogenesis pathway

To investigate the physiological significance of the TmaT-AftD interaction, the AlphaFold 3 structural model was used to generate AftD variants with several amino acid substitutions at the predicted interface. Residues L397, L398, W411, R414, and Q999 at the predicted interface were all changed to alanine **(Fig. 8A).** The resulting AftD variant (AftD^IV^) accumulated in cells to levels equivalent to AftD (WT) but failed to interact with TmaT (**Fig. 8B-C and S8**). Furthermore, AftD^IV^ failed to complement an *aftD* deletion. The cells were susceptible to AU1235 and had an altered trehalose mycolate profile **(Fig. 8B-C).** Production of AftD^IV^ in Δ*aftD* cells also failed to complement the altered 2-AzFPA labeling phenotype **(Fig. 8D).** Based on these results, we conclude that the TmaT-AftD interaction is required for mycomembrane biogenesis via the CmpL4 pathway and for normal arabinan biogenesis.

**Figure 8.**
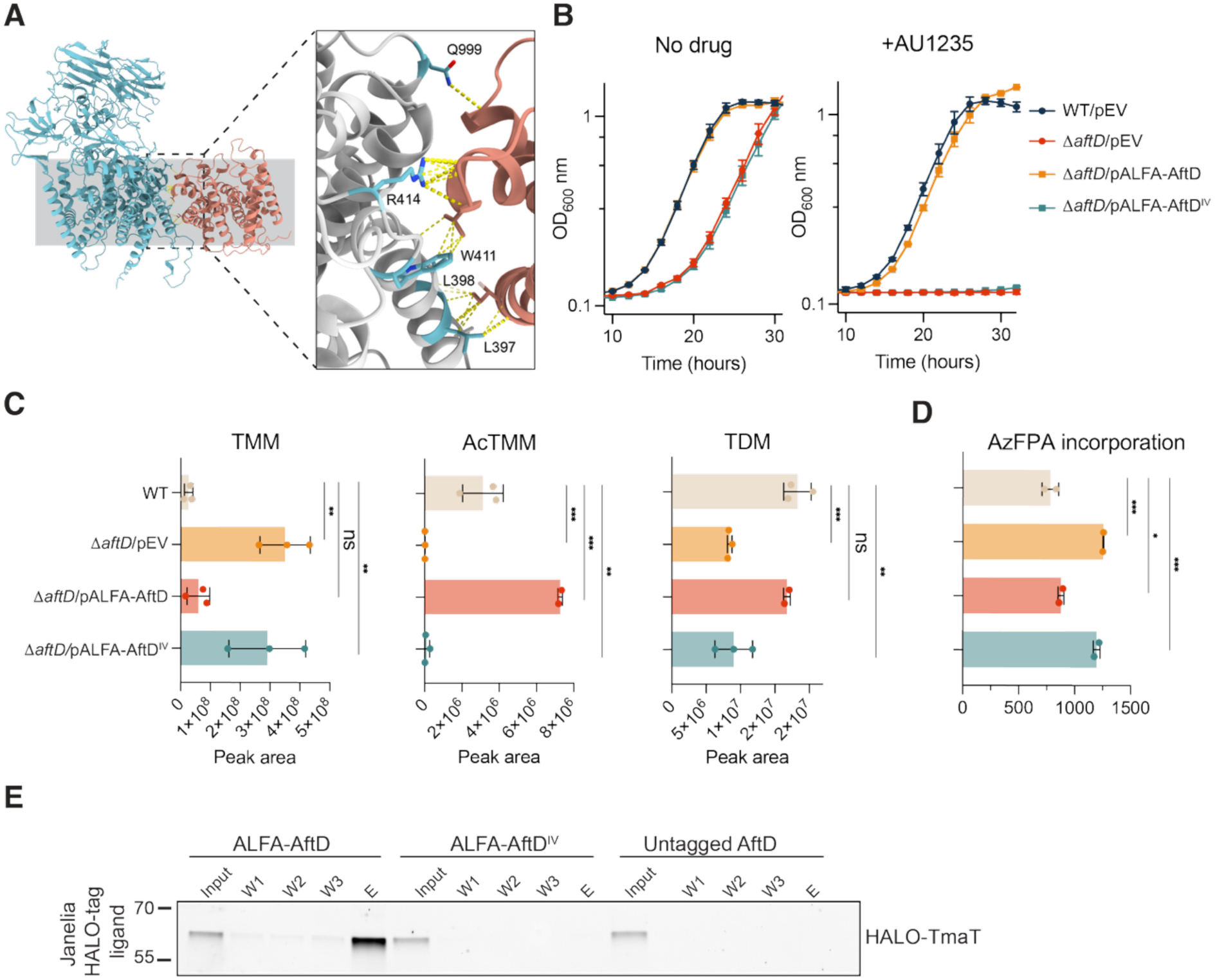
Disruption of the TmaT-AftD interaction impairs mycomembrane assembly. **A**) AlphaFold 3 predicted structure of the TmaT-AftD complex highlighting the interaction interface (left), with a zoomed view of key AftD-interface residues in teal (right)^50^. Residues L397, L398, W411, R414, and Q999 are shown as sticks with predicted contacts to TmaT indicated by yellow dashed lines. **B)** Cultures of strains H2202 [MB001/pEV] H5952 [Δ*aftD*/pEV)], H5768 [Δ*aftD*/pALFA-AftD], and H5874 [Δ*aftD*/pALFA-AftD^IV^] were grown in the presence or absence of AU1235 and growth monitored as in **2C**. **C)** LC-MS Quantification of peak areas for the indicated lipids extracted from the strains in (**B**) as in **2E**. Statistical comparisons were performed using one-way ANOVA with each strain compared to WT (** p<0.01, *** p<0.001, ns= not significant). Data represent individual biological replicates with mean shown. **D)** Flow cytometry analysis of 2-AzFPA (150μM) labeling of the stains in (B) as in **6B**. **E)** Co-immunoprecipitation assay pulling on ALFA-AftD [H5534], ALFA-AftD^IV^ (interface variant) [H5533], or untagged AftD [H5878] using ALFA-resin and probing for HALO-TmaT. Input, wash fractions (W1-W3), and elution (E) were treated with Janelia HALO-tag ligand (Fluor 549) and resolved by SDS-PAGE and imaged on a Typhoon FLA 9500 imager with a TAMRA 532nm filter.

## DISCUSSION

The acetyltransferase TmaT has been known to play a role in mycomembrane assembly for some time. However, in *Cglu* where it was first discovered, it has remained unclear whether TmaT is required for the normal function of both CmpL mycolate transporters or if it specifically functions in one of the two transport pathways. To differentiate between these possibilities, we assessed the growth and mycomembrane assembly phenotypes of double mutants that combined a *tmaT* deletion with the inactivation of CmpL1 or CmpL4. The Δ*tmaT* Δ*cmpL1* double mutant was found to phenocopy cells inactivated for both CmpL1 and CmpL4 whereas the Δ*tmaT* Δ*cmpL4* mutant had a phenotype equivalent to the Δ*tmaT* single mutant. The genetic results therefore indicate that TmaT is required for the activity of the CmpL4 mycolate transport pathway but not for CmpL1 function. Additional genes encoded in the *tmaT* locus were also found to be required for mycomembrane assembly via the CmpL4 pathway, including the arabinosyltransferase AftD. AlphaFold modeling and co-purification experiments indicate that TmaT and AftD interact, and our genetic results suggest that TmaT and AftD functions are mutually interdependent. Our findings thus implicate the TmaT-AftD complex in mycomembrane assembly, providing a potential means by which the process may be coordinated with AG biogenesis. An independent study in *Msmeg* has also demonstrated a role for the TmaT-AftD complex in mycomembrane and AG biogenesis, indicating that this interaction and its role in envelope biogenesis is conserved^24^.

### Potential differences in substrate preference between CmpL1 and CmpL4

Our finding that TmaT is specifically required for the function of the CmpL4 pathway suggests that the CmpL4 transporter requires ^Ac^TMM as a substrate whereas CmpL1 does not. Thus, in the absence of TmaT, it is likely that most of the mycolate transport flux moves through CmpL1. Whether or not CmpL1 activity is positively or negatively affected by TMM acetylation requires further investigation. Although an ^Ac^TMM species analogous to that observed in *Cglu* has not been detected in mycobacteria, the essentiality of TmaT in these organisms suggests that the MmpL3 transport pathway also requires a modified TMM molecule to function properly. Thus, our results suggest that CmpL4 is the functional corynebacterial equivalent of MmpL3 in mycobacteria despite CmpL1 being targeted by MmpL3 inhibitors and sharing greater sequence similarity with MmpL3.

### Topology of TmaT activity and its role in mycolic acid transport

TmaT has previously been proposed to acetylate TMM on the cytoplasmic side of the membrane to facilitate its transport to the exterior face of the membrane by MmpL/CmpL transporters^4^. However, TmaT belongs to the AT3 family of acetyltransferases (InterPro: IPR002656) that typically function by acetylating their substrates on the extra-cytoplasmic side of the membrane^51–54^. This reaction involves the binding of an acetyl-CoA or related acyl donor through a cytosolic opening in the acyltransferase, positioning the acetyl group for transfer to a substrate on the opposite side of the membrane. Accordingly, an AlphaFold model of TmaT with a bound acetyl-CoA molecule has a cofactor orientation resembling its positioning in solved structures of other AT3 family proteins in complex with their acyl donor (**Fig. S9**)^55,56^. Although further structural and biochemical analyses are required to validate an extra-cytoplasmic activity for TmaT, this reaction topology seems more likely than the previously proposed model for the cytoplasmic modification of TMM. Assuming TmaT functions at the membrane surface like other AT3 family proteins, the role of TMM acetylation in its transport requires reevaluation. If it is not promoting the flipping of TMM across the membrane, what is the purpose of the modification? One attractive possibility is that it marks a subpopulation of TMM molecules for a specific “destination” (protein-linked, AG-linked, or TDM production) following extraction from the membrane by the MmpL3/CmpL4 transporters. Testing this and other potential roles for TmaT in TMM transport will shed additional light on the molecular mechanisms underlying mycomembrane biogenesis.

### Role of AftD and other factors encoded in the tmaT locus

AftD was originally identified as an AraT enzyme with ⍺1-3 branching activity that participates in the biogenesis of the arabinan component of AG and lipoglycan polymers^47^. An alternative activity was proposed in a subsequent study in which an ⍺1-5 transferase activity for AftD was detected, suggesting that it plays a role in extending the arabinan branches of AG^46^. Regardless of the precise role of AftD in AG biogenesis, its requirement for TmaT function places it at the nexus of at least two and possibly three critical envelope biogenesis pathways (AG, lipoglycans, and the mycomembrane). Whether TmaT also affects AftD activity is unclear, but the observation that TmaT inactivation phenocopies the effect of AftD inactivation with respect to arabinan label incorporation suggests that the dependency is mutual. Notably, both mutants show an increase in 2-AzFPA labeling. Although a defect associated with an arabinosyltransferase pathway might be expected to reduce arabinan labeling, the observed increase suggests that loss of TmaT or AftD alters the process, for example, by increasing probe accessibility or dysregulated activity of other Aft proteins. How the two enzymes might influence each other’s activity remains to be determined. It is possible that each protein allosterically stimulates the activity of the other. Alternatively, the product of one enzyme may be needed for the activity of the other and vice versa. Given that formation of the AftD-TmaT complex is required for normal envelope biogenesis, we favor the former possibility, but other more complicated models cannot be excluded at this time.

Several factors encoded in the conserved *tmaT* locus were previously shown to be required for ^Ac^TMM formation^29,30^ and found here to function in the CmpL4 transport pathway along with AftD and TmaT. Their roles in mycomembrane biogenesis also require clarification. Unlike AftD, AlphaFold does not predict direct 1-1 interactions between these factors and TmaT, but it remains possible that a higher-order complex involving multiple components encoded in the *tmaT* locus is involved in promoting TMM transport via CmpL4 in corynebacteria and by MmpL3 in mycobacteria. Notably, Foldseek^57^ searches of the predicted structures of MmpA and Cgp_3166 indicate that they may be glycosyltransferases, suggesting their involvement in the synthesis and/or modification of one or more envelope glycans. Additionally, Cashmore and colleagues noted that the *Cglu* and *Mtb* MmpA shares partial homology with the first 11 transmembrane domains of *Cglu* or *Mtb* AftD, respectively.^20^ Therefore, like AftD, complexes involving these proteins and TmaT and/or other potential partners may provide additional links between mycomembrane assembly and the biogenesis of the underlying envelope layers. Further characterization of these potential connections promises to reveal at least some of the mechanisms corynebacteria and mycobacteria use to build their complex surface architecture in a uniform and coordinated fashion.

## METHODS AND MATERIALS

### Bacterial Strains and Growth Conditions

All bacterial strains used in this study are listed in **Table S2**. Unless otherwise specified, experiments were conducted using *Corynebacterium glutamicum* MB001, a prophage-free derivative of ATCC13032^58^. *C. glutamicum* strains were cultured in BD BBL Brain Heart Infusion Broth (BHI) medium (cat# BD211059) or BHI supplemented with 9.1% sorbitol (BHIS) at 30°C with aeration, except when strains carried the temperature-sensitive plasmids pEWL89 (Cre recombinase) or pEWL103 (recombineering vector), which were maintained at 25°C^59^. Agar was made with the addition of 1.5% BD BACTO agar (w/v) (cat# 214040) to BHI, BHIS, or LB. Competency medium is described below.

Ectopic expression was induced only for co-immunoprecipitation experiments with 0.5 mM theophylline (*riboE1*-integrated constructs)^60,61^, or 50 µM IPTG for *Ptac*-driven plasmids derived from pTGR5^59,62^. *Escherichia coli* DH5α (λpir) and NEB10-β strains used for cloning were grown in LB medium (1% tryptone, 0.5% yeast extract, 0.5% NaCl) at 37°C with aeration, except when carrying pCRD206 allelic-exchange derivatives, which were cultured at 30°C.

When appropriate, *C. glutamicum* cultures were supplemented with the following antibiotics: kanamycin (15 µg/mL), chloramphenicol (3.5 µg/mL), or apramycin (12.5 µg/mL). *E. coli* strains were grown with kanamycin (25 µg/mL), chloramphenicol (25 µg/mL), or apramycin (50 µg/mL) for plasmid maintenance and cloning.

### Growth curves

*C. glutamicum* cells were inoculated from single colonies into 5mL of BHI and grown overnight at 30°C on a roller drum (terminal OD_600_ ∼5-6). Overnights were diluted 1:50,000 into fresh BHI containing either DMSO or freshly resuspended 25 µM AU1235 (from a 10 mM stock in 100% DMSO). Cells were gently vortexed to mix and 200 µl samples were aliquoted in technical triplicate into a 96-well Corning cell culture cluster flat bottom polystyrene plate. The lid was parafilmed to the base plate with two layers. The culture OD_600_ was tracked in either a Versa max microplate reader using continuous orbital shaking at 30°C or in an Aglient BioTek Gen5 with double orbital continuous shaking at 30°C. Measurements were plotted in GraphPad Prism version 11.

### Agar growth assays

*C. glutamicum* cells were inoculated from single colonies into 5 mL of BHI and grown overnight at 30°C on a roller drum. Overnights were standardized to an OD_600_ of 0.6 and then diluted 1:10 into BHI in a 96-well Corning cell culture cluster flat bottom polystyrene plate with a final volume of 200 µL. Cells were serially diluted 1:10 down to a 10^−6^ dilution. Four microliters of each dilution were then spotted onto a BHI plate, dried, and plates were incubated at 30°C overnight. Plates were imaged using a Nikon Z5 II Mirrorless camera with a NIKKOR Z MC 50mm f/2.8 Macro Lens.

### Plasmid Construction

Primers for cloning of plasmids were purchased from IDT and are listed in **Table S3**. PCR for cloning was conducted using NEB Q5 2X Mastermix High-Fidelity DNA polymerase (cat# M0492). Plasmids were assembled via isothermal assembly (ITA). For site-specific mutagenesis, plasmids were amplified with NEB Q5 polymerase or Roche KAPA HiFi HotStart ReadyMix (cat# 07958927001) and either treated with NEB KLD enzyme mix (cat# M0554S) or ITA. DNA was transformed into NEB10-beta or DH5α (λpir) *E. coli* cloning strains by electroporation or heat shock at 42°C for 45 seconds, respectively. All plasmids used in this study are listed in **Table S4**. A special note for cloning of *aftD*, an extra 84 bases upstream of the putative ATG were included in the construction of *aftD* complementation vectors.

### Strain Construction

*C. glutamicum* competent cells were prepared as described previously^63,64^. Briefly, 5 ml overnight cultures grown in BHIS were subcultured in competency medium consisting of BHI supplemented with 91 g/L sorbitol, 0.1% Tween 80, 0.4 g/L isoniazid, and 25 g/L glycine at a 1:50 or 1:100 dilution and grown at 30°C for 3 hours or at 18°C overnight to mid-logarithmic phase. Cultures were then washed three times with cold 10% glycerol by centrifugation at 4,000 x g for 10 minutes. Final cell density was adjusted to an OD_600_ of approximately 20. DNA (50-500ng) was added to cells, transferred to a 1 mm electroporation cuvette, and cells were electroporated using the settings 1.8 kV, 25 µFD, 200 ohms. Cells were immediately heat shocked for 6 minutes at 42°C, then incubated for 1-2 hours at 30°C unless a temperature sensitive plasmid was transformed, in which case they were incubated at 25°C for 3 hours.

Gene deletion was performed in one of two ways; Allelic exchange via *sacB* counterselection with pCRD206 as published previously or by recombineering as described in a previous report^59,65^. Briefly, *C. glutamicum* was transformed with pEWL103 and a saturated culture was grown in BHIS containing apramycin (12.5ug/ml) at 25°C overnight. Cells were diluted 1:100 and grown in transformation media at 25°C for 4 hours. The SSAP/SSB pair was then induced by adding IPTG and theophylline to final concentrations of 1 mM each and growth was continued for another 4 hours at 25°C. Competent cells were then prepared by washing with 10% glycerol as described above. Linear dsDNA cassettes were constructed using a pair of 70-mer oligonucleotides. The 70-mer oligonucleotides were used to amplify the kanamycin resistance cassette flanked by LoxP66 and LoxP71 sites located on pEWL74. Each 70-mer oligonucleotide contained 20 bp of homology to the cassette and 50 bp of homology to the genomic region of interest. Induced competent cells carrying pEWL103 were transformed with 500 ng of purified linear dsDNA following the conditions described above. Cells were recovered for 2 hours at 30°C, and the entire transformation was plated on BHIS agar containing kanamycin to select for recombinants. Recombinants were confirmed using colony PCR. Plating the transformation at 30°C was sufficient curing pEWL103, where appropriate^59^. Electrocompetent cells of recombineered strains were prepared as described above. Plasmid pEWL89 was used to remove the kanamycin cassettes via Cre recombination^59^. It was transformed by electroporation. Cells were recovered and plated at 25°C on BHIS apramycin. Transformants were streaked on BHIS at 30°C and then patched to ensure that the kanamycin resistance cassette had been removed. In the case of pEWL89, patching at 30°C was sufficient to cure the plasmid. Excision of the kanamycin resistance cassette was confirmed by colony PCR.

### Phenotypic correlation profiles with Tn-seq data

Raw sequencing data were obtained from the Sequence Read Archive (SRA), accession number PRJNA610521^36^. Reads were aligned to the *Corynebacterium glutamicum* MB001 genome (NCBI accession number CP005959) using the local alignment option of bowtie2 (v2.5.4)^66^. Aligned reads were sorted and filtered for a minimum mapping quality of 40 with samtools (v1.22.1) view^67^. Sequencing depths per coordinate were obtained for each strand with samtools depth. Transposon insertion counts were quantified for every coordinate as the change in sequencing depth between the coordinate and its direct neighbor using a custom R (v4.1.1) script. Read counts were subsequently summed for every genetic element annotated with a locus_tag. These gene-wise summed read counts were normalized using the variance stabilizing transformation (vst) of R package DESeq2 (v1.46.0)^68^. Using these fitness scores, Pearson correlation coefficients were computed across all libraries, except the input libraries (1g_A, 1g_B and 1g_C), for every pair of genes with non-zero cross-library variance.

### 6-TMR-Trehalose mycomembrane labeling assays

*C. glutamicum* cells were inoculated from single colonies into 5mL of BHI and grown overnight at 30°C on a roller drum. Cells were subcultured 1:50 into BHI until they reached OD_600_ of ∼ 0.4-0.6. An equivalent of OD_600_ = 0.3 was pelleted and resuspended in 1X PBS (137mM NaCl, 2.7mM HCl, 8mM Na_2_HPO_4_, 2mM KH_2_PO_4_ pH 7.4) supplemented with a final concentration of 100 µM 6-TMR-Trehalose (10 mM stock in 100% DMSO; Tocris Bioscience cat# 6802). Cells were incubated for 1 hour in the dark at room temperature in a final volume of 100 µl. Samples were then washed twice in 1X PBS with centrifugation at 10,000xg for 5 minutes between washes and a final resuspension of 1 mL in 1X PBS. To a black-walled clear bottom 96 well plate (Costar® Assay plate cat# 3631) 200 µL of each sample was aliquoted in technical triplicate, including a PBS only control. The OD_600_ of each sample was measured and 6-TMR-Trehalose incorporation was measured at an excitation of 532 nm and emission of 580 nm on a Tecan Infinite M Plex. Incorporation is reported as 6-TMR-Trehalose/OD_600_ with calculations performed in Microsoft Excel and graphs generated using GraphPad Prism version 11. These experiments were repeated at least three independent times.

### Co-Immunoprecipitations

Single colonies from strains H#5534, H#5533, and H#5878 were inoculated into 10 ml each of BHI containing 3.5 µg/ml of chloramphenicol and grown on a roller drum at 30°C overnight. The next morning cells were subcultured into 2L flasks containing BHI supplemented with 3.5 µg/ml chloramphenicol, 500 µM theophylline, and 50 µM IPTG. Cells were grown shaking at 250 rpm at 30°C until OD_600_ of ∼3-3.5 was reached. Cells were pelleted, supernatant was removed, and pellets were flash frozen and stored at −80°C until needed.

To conduct the Co-IP, pellets were thawed to room temperature and resuspended in 25 mL of lysis buffer (1X PBS pH 7.4, Roche cOmplete™ protease inhibitor cocktail tablets, 2 µg/mL lysozyme, and 100kU of Pierce™ Universal nuclease (cat#88702)). Cell lysate was rotated head over tail at 30°C for 1 hour to allow lysozyme lysis. Lysates were then passaged four times through a Constant Systems continuous F1 cell disruptor at 40,000 psi. Large cell debris and intact cells were pelleted at 10,000xg at 4°C for 10 minutes, and supernatant was transferred to 70 mL polycarbonate tubes (38×102mm; cat #: 355622) and centrifuged at 100,000xg (35,000xrpm) in a Type 45 Ti fix-angled rotor in a Beckman Optima XE ultracentrifuge for 1 hour. Supernatant was removed and the membrane pellet was resuspended in 3 mL membrane solubilization buffer (1X PBS pH 7.4, 1% n-dodecyl β-D-maltoside (DDM)) by douncing, then tumbled head over tail for 1 hour at 4°C. The membrane fraction was diluted to 0.1% DDM with the addition of 1X PBS pH 7.4. Input samples were removed and saved for later. Equilibrated ALFA-selector PE magnetic resin was added (40 µL of 50% slurry per sample), and solubilized membrane proteins were incubated with resin for 15 minutes at 4°C rotating end over end was performed. After incubation, samples were centrifuged at 100xg for 1 minute to gently pellet resin. All supernatant except 100μL was removed and resin was resuspended and transferred to 1.7mL tubes. Magnetic resin was washed 4 times with the following buffers: Wash 1 and Wash 4 (1X PBS pH 7.4 (equivalent to 137mM NaCl), 0.1% DDM), Wash 2 and Wash 3 (1X PBS pH 7.4 plus an extra 63mM NaCl to get 200mM NaCl, 0.1% DDM). Washes were incubated for 5 minutes each rotating at 4°C. Proteins were eluted from the resin by incubation at room temperature for 20 minutes while rotating in Elution buffer (1X PBS pH 7.4, 200 µM ALFA elution peptide, 0.05% DDM). Input, wash, and elution fractions were mixed with 6X SDS-PAGE reducing loading dye (final concentration of 62.5mM Tris-HCl pH 6.8, 1.5% SDS (w/v), 8.3% glycerol, 0.0125% bromophenol blue, 1.5% 2-mercaptoethanol) and resolved for 45 minutes at 200V on Biorad 4-20% Mini-PROTEAN^®^ TGX™ Precast protein gel using 1X Tris-Glycine-SDS running buffer (250mM Tris-HCl, 1.92M glycine, 1% SDS). The ladder used for all SDS-PAGE was PageRuler™ Plus Prestained protein ladder 10 to 250kDa (cat#26620).

### HALO-Ligand and Immunoblot Analysis

To assess Co-elution of Halo-tagged AftD or Halo-tagged TmaT from Co-immunoprecipitation assays, elution samples were incubated with 200 nM Janelia Fluor 549 HaloTag Ligand (Promega cat# GA1111) for 5 minutes prior to SDS-PAGE analysis. Gels were imaged on a Typhoon FLA 9500 using TAMRA 532nm (LPG) filter. For Western blot analysis of ALFA-tagged proteins, resolved precast gels were transferred using a Biorad semi-dry Trans-Blot Turbo transfer system using 1X Biorad Trans-Blot Turbo buffer, Biorad Trans-blot Turbo midi filter paper (cat#12023956) and Biorad Immun-Blot PVDF membrane (cat1620177) hydrated in 100% ethanol. System transfer settings were 2.5A, 25V for 7 minutes. Transferred PVDF membranes were incubated in 1X PBST containing 5% milk (10mM anhydrous Na_2_HPO_4_, 137 mM NaCl, 2.7mM KCl, anhydrous 1.5mM KH_2_PO_4_, 1% Tween 20, pH 7.4) for 30 minutes at room temperature with gentle rocking. To probe for ALFA-tag, 1X PBST+5% milk containing 1:5000 dilution of sdAb anti-ALFA HRP antibody (NanoTag) was incubated on the blot for 1 hour at room temperature or overnight at 4°C. The blot was washed 3 times with 1X PBST and developed with SuperSignal™ West Pico PLUS Chemiluminescent substrate (cat#34580) and imaged on a Biorad ChemiDoc™ MP imaging system using chemiluminescence filter.

### Co-purification of His_10_-TmaT and FLAG-AftD from *E. coli*

*E. coli* C43 chemically competent cells were transformed with pES113 and plated on LB-Carb100 at 37°C. 5-10 colonies were then inoculated into 50 mL of terrific broth supplemented with 100 μg/mL carbenicillin and grown at 30°C overnight. Overnight cultures were then diluted 1:100 in terrific broth and grown at 37°C until an OD_600_ between 0.8-1.0 was reached. Protein production was induced with 500 μM IPTG, and cultures were grown overnight at 23°C with aeration. Cells were then pelleted, washed in 1X PBS, and frozen at −80°C until further use. Thawed cell pellets were then resuspended in lysis buffer (50 mM HEPES pH 7.4, 400 mM NaCl, 10% (v/v) glycerol, 1 mg/mL lysozyme, and 1:50,000 (v/v) Benzonase). Cells were then lysed on the EmulsiFlex-C5 cell disruptor (Avestin) at 15,000 psi for 5-6 cycles. Lysates were centrifuged at 10,000xg for 10 minutes to remove cell debris and unlysed cells. The supernatant was transferred to Beckman ultracentrifuge tubes and ultracentrifuged at 140,000 xg for 45 minutes at 4°C. The supernatant was discarded, and the membrane pellet was homogenized with a Dounce homogenizer in 6 mL of Homogenization buffer (50 mM HEPES pH 7.4, 400 mM NaCl, 10% (v/v) glycerol, and 1% (m/v) DDM) and diluted with homogenization buffer to a final volume of 40 mL. The resulting homogenate was rotated end over end at 4°C for 1.5 hours. A second ultracentrifugation step was carried out to remove non-homogenized/solubilized membranes. The resulting clarified membrane fraction was batch bound with 1 mL of pre-equilibrated TALON resin for 30 minutes at 4°C with 1 mM of imidazole. The resin was washed three times with 20 column volumes each of Buffer A with varying concentrations of DDM and imidazole (first wash: 1% m/v DDM and 2 mM imidazole, second wash: 0.2% m/v DDM and 4 mM imidazole, third wash: 0.1% m/v DDM and 6 mM imidazole). 10 column volumes of elution buffer (Buffer A with 150 mM imidazole and 0.05% m/v DDM) was then added to the column, and the resin was rocked in the elution buffer for 10 minutes. The elution was concentrated on a 100 kDa molecular weight-cut off Amicon column at 4°C down to 1 mL. The concentrated elution was then further purified by size exclusion chromatography on a Superdex 200 10/300 column (Cytiva) in SEC Buffer (50 mM HEPES pH 7.4, 150 mM NaCl, 5% (v/v) glycerol, and 0.05% (m/v) DDM). The SEC fractions containing pure complex were pooled and concentrated to 22 µM as determined by absorbance at A280.

### AlphaFold modeling of protein complexes

AlphaFold 3.0 was used to predict structures^50^. Protein sequences were from the MB001 genome into AF3. PAE plots were visualized using Predictomes.org to obtain plots shown in this manuscript for figure S7, and all remaining PAE plots were visualized in PAE-Viewer^69^. Models were visualized in PyMOL (Schrodinger) or ChimeraX.

### AzFPA Flow Cytometry

Experiments were performed following as described previously with modifications^49^. In brief, *C. glutamicum* strains were grown to saturation overnight in BHI with the reported antibiotic concentrations. Cultures were then normalized to an OD_600_ of 0.2 in BHI with the reported antibiotic concentrations and seeded in a Corning black 96-well plate (100 μL). 2-AzFPA was added to a final concentration of 150 μM. Cultures were grown shaking for 4 h at 30 °C. Cells were pelleted by centrifugation for 5 min at 3000 x g. The pellets were washed with ice-cold phosphate buffered saline (PBS) supplemented with 0.025% Tween 80 (100 μL) twice. Cells were then resuspended in PBS and AFDye™ 647 DBCO (Lumiprobe # 2G8F0) was added to a final concentration of 150 μM. The samples were stained for 1.5 h rotating at 30 °C. The stained cells were pelleted for 5 min at 3000 x g. The supernatant was removed, and the pellet was washed with PBS supplemented with 0.025% Tween 80 and 0.5% (w/v) BSA once, then PBS supplemented with 0.025% Tween 80 once. The final pellet was then resuspended in 1 mL PBS and analyzed using a BD FACS Symphony A1 Analyzer. A total of 100,000 cells were counted at the low flow rate. Flow cytometry analysis was performed in triplicate, and representative scatter plots are shown (**Figure S5**). The dye only controls were analyzed first to set gates. Data were analyzed using the FlowJo software package (FlowJo LLC). Mean fluorescence intensity was calculated using a geometric mean.

### Lipid extraction

*Cell growth.* Extraction of non-covalently bound lipids was performed using a modified Bligh and Dyer method as follows: Single colonies from BHIS plates were inoculated into 2 or 5ml of BHI (in biological triplicate or quadruplicate) in 18×150mm glass culture tubes and rotated on a roller drum overnight at 30°C. The next morning cells were subcultured 1:50 into 5ml of BHI containing appropriate antibiotics (see figure legends for details per experiment). Cells were grown rotating on a roller drum at 30°C until OD_600_ of 3-5 was reached. Cells totaling an OD_600_ of 12 were pelleted at 4,000xg for 5 minutes, the supernatant was removed, and cells were flash frozen at −80°C until extraction.

*Extraction.* Cells were thawed at room temperature, washed with 500 µl of 1X PBS pH 7.4, and transferred to PYREX 10 ml glass conical centrifuge tubes (cat# 99502-10). Cells were pelleted at 3,000xg for 10 minutes, the 1X PBS supernatant was removed, and pellets were resuspended in 1:2 chloroform:methanol in a final volume of 1.2 ml, where chloroform and methanol were added consecutively. Samples were rocked at room temperature for 1 hour. Samples were centrifuged at 3,000xg for 10 minutes and the supernatant was transferred to 2 ml glass HPLC vials to be dried by nitrogen gas stream in the fume hood. The remaining pellet was resuspended in 1:1 chloroform:methanol in a final volume of 1.2 ml, added consecutively, and were rocked at room temperature for 1 hour. Samples were centrifuged at 3,000xg for 10 minutes and the supernatant was pooled with previous supernatant in 1 ml HPLC vials and dried by nitrogen gas stream. The remaining pellet was resuspended in 2:1 chloroform:methanol and rocked for 1 hour. Samples were then centrifuged at 3,000xg for 10 minutes and the supernatant was added to 2ml HPLC vials with other supernatant fractions and dried down by nitrogen. Lipid samples were resuspended in 600 µl of chloroform, and 600 µl of water was added. Samples were vortex vigorously and centrifuged at 3,000xg for 10 minutes. The organic bottom layer was removed and added to fresh 2ml HPLC vials, where samples were dried by nitrogen stream. Samples were resuspended in a final volume of 600 µl 1:2 chloroform:methanol to be injected for LC-MS analysis.

### LC-MS Analysis

Lipid extract samples were analyzed by liquid chromatography high resolution mass spectrometry using a Vanquish – Orbitrap Exploris 240 system (Thermo Electron, Madison, WI). Samples were injected (5 μL) into an Acquity Premier HSS T3 1.8 μm, C18 column (2.1×100mm), with guard, using 6:4 acetonitrile:water with 10 mM ammonium formate and 0.1% formic acid (A) and 9:1 isopropyl alcohol:acetonitrile with 10 mM ammonium formate and 0.1% formic acid (B) at 0.3 ml/min and 45 °C. Analytes were separated using a gradient from 30 to 98% B over 10 minutes, followed by a 98% B wash and equilibration to 30% B. Mycolates were detected in electrospray positive ion mode as [M+NH^4^]^+^: TMM 838.6255, AcTMM 880.6362, TDM 1317.0966). Peak areas were calculated using Thermo Xcalibur 4.5.445.18 Quan Browser. LC-MS/MS was performed and described in the supplemental figures 1-3 to confirm structures in the absence of standards.

## ACKNOWLEDGEMENTS

The authors would like to thank members of the Bernhardt, Walker, Rudner labs, and the members of the *Corynebacterium* subgroup for excellent advice and many helpful discussions. We also acknowledge the Analytical Chemistry Core at Harvard Medical School for access to LC-MS instrumentation and their expertise. This work was supported by Investigator funds from the Howard Hughes Medical Institute (to T.G.B), the National Institutes of Health F32 1F32AI191488-01 (to A.R.P.), the Helen Hay Whitney Foundation fellowship (to V.M.M), the Swiss National Science Foundation Postdoc Mobility fellowship P500PB_225439 (to V.d.B.), the National Institutes of Health R01 AI148752 and U19 AI158028 (to S. W.), and R01 AI-126592 (to L.L.K.).

## Author Contributions

A.R.P., E. D. S., V. M. M., M. J., S. W., and T. G. B. designed research; A. R. P., E. D. S., V. M. M. performed research; V. d. B. analyzed data; L.L.K. provided chemical probes; A. R. P., E. D. S., V. M. M., S. W., and T. G. B. wrote the paper.

## SUPPLEMENTAL FIGURES

**Figure S1.**
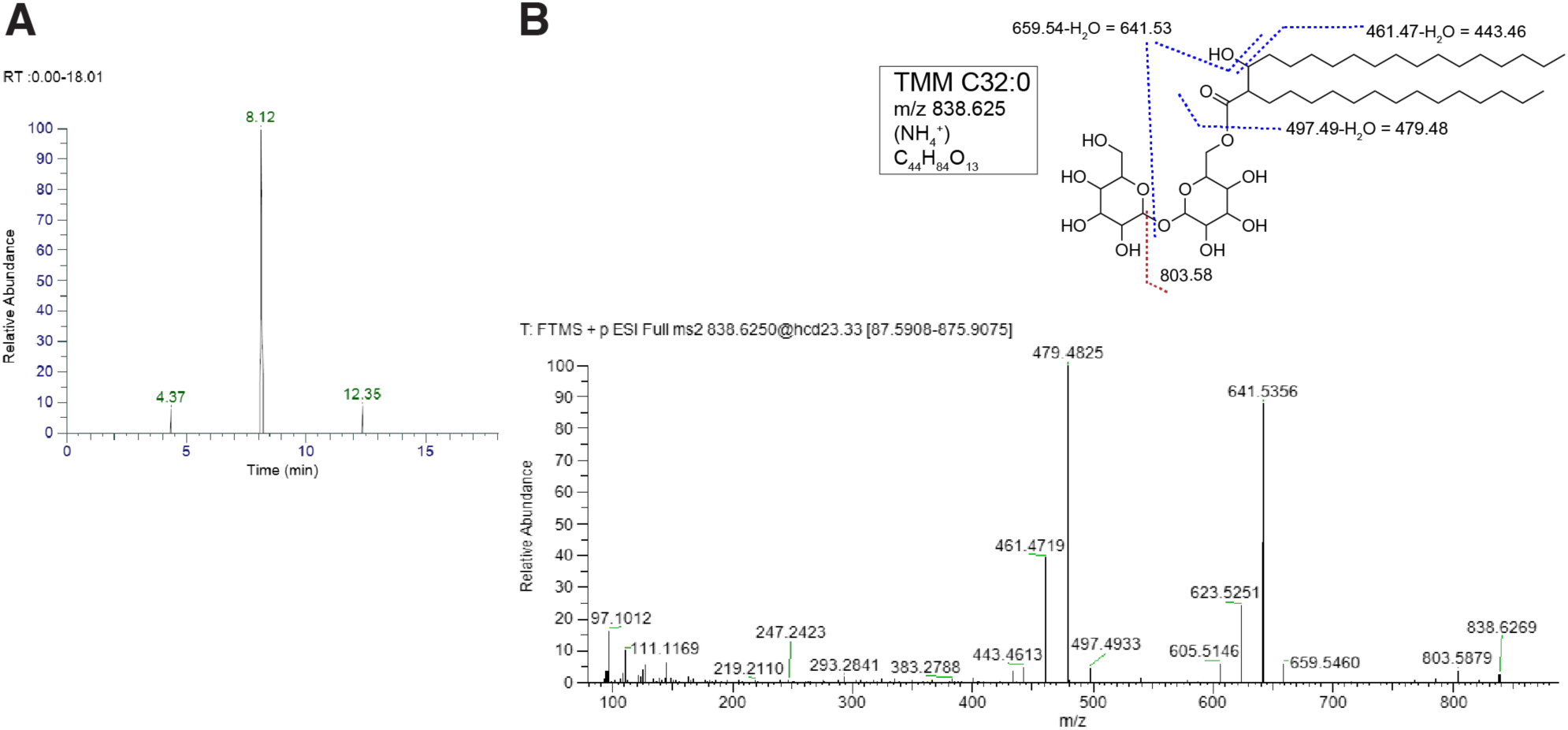
Structural characterization of trehalose monomycolate (TMM) C32:0 from *Corynebacterium glutamicum* MB001 by high-resolution LC-MS/MS. **A)** Representative MS2 spectrum of TMM C32:0 detected as [M+NH₄]⁺ at *m/z* 838.625 (C_44_H_84_O_13_, HCD 23.33 eV) from *C. glutamicum* MB001 grown in BHI medium. Chromatographic retention time 8.12 min. **B)** Annotated structure of TMM C32:0 showing diagnostic fragment ion assignments. Blue dashed lines indicate cleavage at the mycolate ester bond (C6-position of trehalose) and mycolic acid chain, generating acylium ions at *m/z* 497.49 (pre-dehydration), 479.48, 461.47, and 443.46 (sequential −H₂O losses). Red dashed lines indicate cleavage at the glycosidic bond between the two glucose units of the trehalose headgroup, generating oxocarbenium ions at *m/z* 659.54, 641.53, 623.53, and 605.51 (sequential −H₂O losses). The fragment at *m/z* 803.58 arises from loss of the non-mycolylated glucose as a neutral, retaining the intact mycolate-bearing glucose-mycolate ion. Low-mass fragments at *m/z* 97.10, 111.12, 219.21, 247.24, 293.28, and 383.28 correspond to aliphatic chain fragments along the mycolic acid α-branch and meromycolate chains. Lipid extraction is described in Materials and Methods. All spectra were acquired on a Thermo Fisher Scientific Orbitrap Exploris 240 mass spectrometer in positive ion mode.

**Figure S2.**
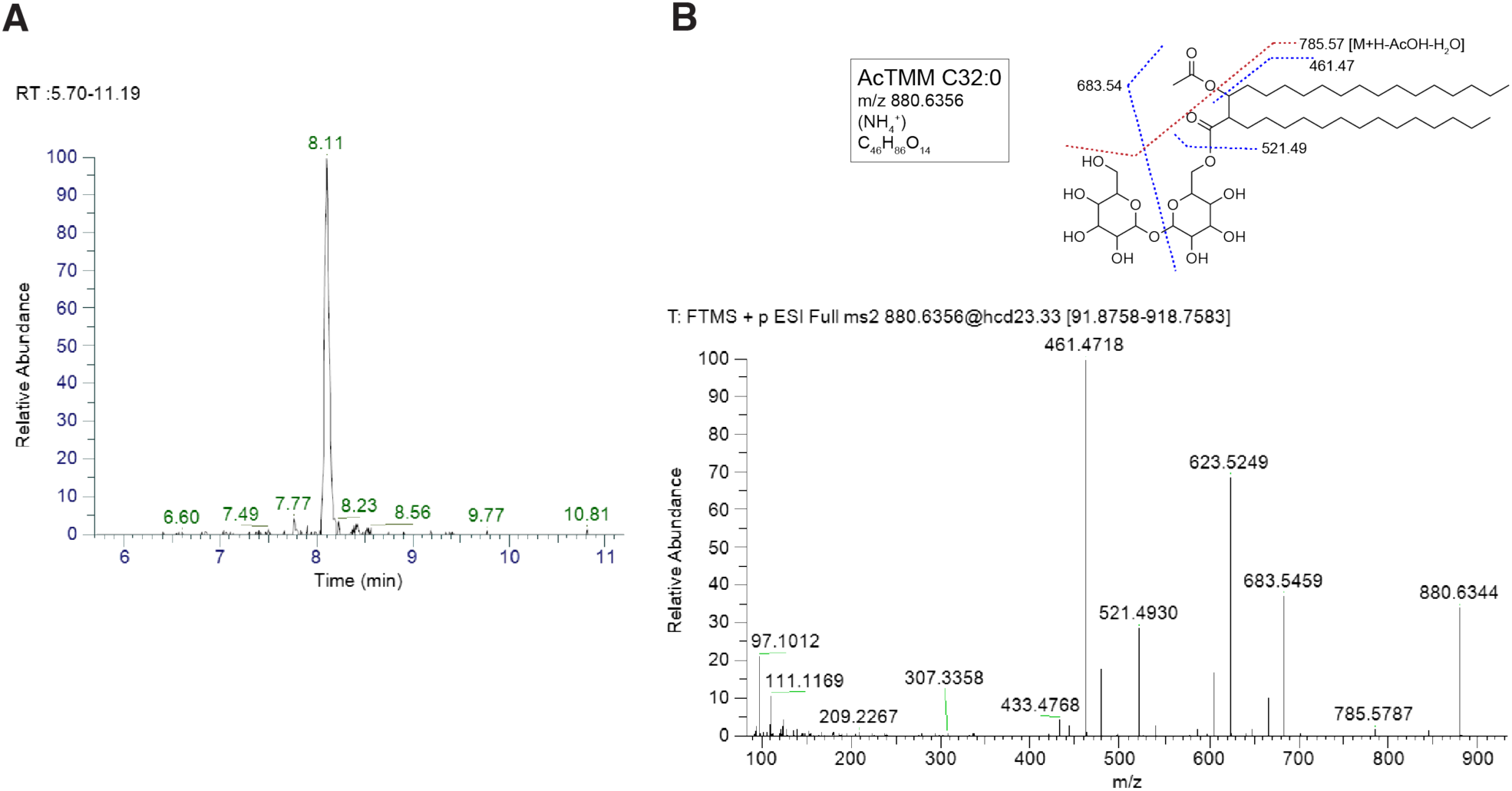
Structural characterization of acetyl-trehalose monomycolate (AcTMM) C32:0 from *Corynebacterium glutamicum* MB001 by high-resolution LC-MS/MS. **A)** Representative extracted ion chromatogram of AcTMM C32:0 detected as [M+NH₄]⁺ at *m/z* 880.636 (C_46_H_86_O_14_, HCD 23.33 eV) from *C. glutamicum* MB001 grown in BHI medium. Chromatographic retention time 8.11 min. **B)** MS2 spectrum and annotated structure of AcTMM C32:0. AcTMM differs from TMM by the presence of an acetyl group on the β-hydroxyl of the mycolic acid chain. Blue dashed lines indicate cleavage at the mycolate ester bond (C6-position of trehalose) and mycolic acid chain, generating acylium ions at *m/z* 521.49 and 461.47. Red dashed lines indicate cleavage at the acetyl ester bond on the β-carbon of the mycolic acid, generating the diagnostic fragment at *m/z* 785.57 ([M+H−AcOH−H₂O]⁺), confirming the presence and position of the acetyl modification, and the glycosidic bond cleavage fragment at *m/z* 683.54. The fragment at *m/z* 623.52 corresponds to the glycosidic bond oxocarbenium ion, shifted −18 Da relative to the analogous TMM fragment (*m/z* 641.53) due to consumption of the β-hydroxyl by acetylation. Low-mass fragments at *m/z* 97.10, 111.12, 209.23, and 307.34 correspond to aliphatic chain fragments along the mycolic acid chains. Lipid extraction is described in Materials and Methods. All spectra were acquired on a Thermo Fisher Scientific Orbitrap Exploris 240 mass spectrometer in positive ion mode.

**Figure S3.**
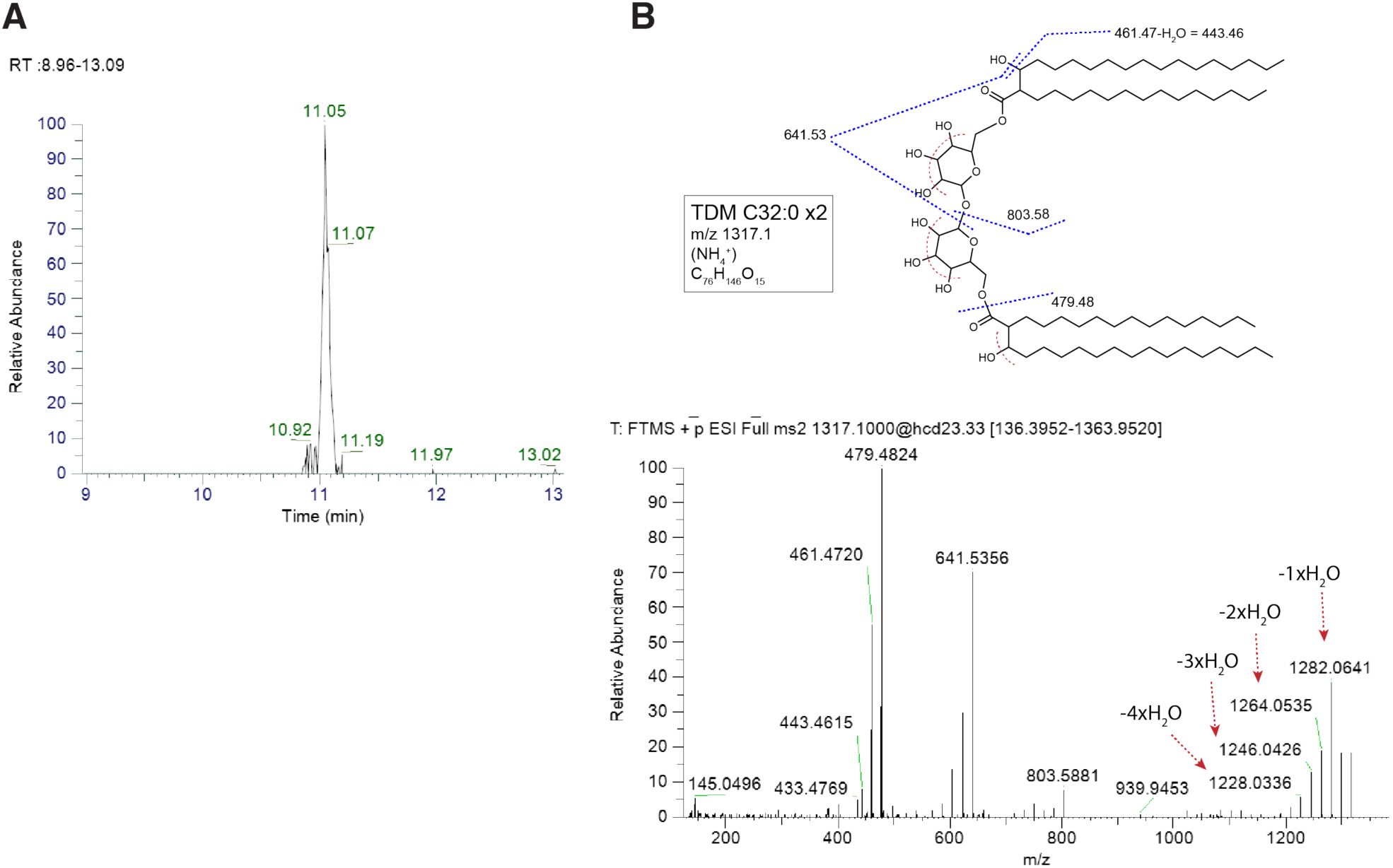
Structural characterization of trehalose dicorynomycolate (TDM) C32:0 x2 from *Corynebacterium glutamicum* MB001 by high-resolution LC-MS/MS. **A)** Representative extracted ion chromatogram of TDM C32:0 ×2 detected as [M+NH₄]⁺ at *m/z* 1317.1 (C_78_H_148_O_15_, HCD 23.33 eV) from *C. glutamicum* MB001 grown in BHI medium. Chromatographic retention time 11.05 min**. B)** MS2 spectrum and annotated structure of TDM C32:0 ×2. Blue dashed lines indicate symmetric cleavage at both mycolate ester bonds (C6 and C6′ positions of trehalose), generating mycolic acid acylium ions at *m/z* 479.48 (base peak) and 461.47, with sequential −H₂O loss giving 443.46. Cleavage at the glycosidic bond generates the oxocarbenium fragment at *m/z* 641.53. The fragment at *m/z* 803.58 arises from loss of one intact mycolic acid chain, generating a TMM-equivalent fragment ion retaining the trehalose headgroup esterified to a single mycolate chain. The fragment at *m/z* 145.05 corresponds to the trehalose oxocarbenium headgroup ion. The fragment at *m/z* 939.95 is assigned as [M+H]⁺ minus the intact trehalose headgroup (−360 Da), retaining both mycolic acid chains as a bis-mycolate fragment ion. Red dashed arrows in the MS2 spectrum indicate sequential neutral losses of H₂O (−18 Da each) from [M+H]⁺ (*m/z* 1300.07), generating a dehydration ladder at *m/z* 1282.06 (−1×H₂O), 1264.05 (−2×H₂O), 1246.04 (−3×H₂O), and 1228.03 (−4×H₂O). The first three losses are attributed to the three free hydroxyl groups of the trehalose headgroup; the fourth loss (−4xH₂O, *m/z* 1228.03) is attributed to the free β-hydroxyl of the mycolic acid chain, consistent with the absence of this fourth loss in AcTMM (Fig. S2), where the β-hydroxyl is consumed by acetylation. Lipid extraction is described in Materials and Methods. All spectra were acquired on a Thermo Fisher Scientific Orbitrap Exploris 240 mass spectrometer in positive ion mode.

**Figure S4.**
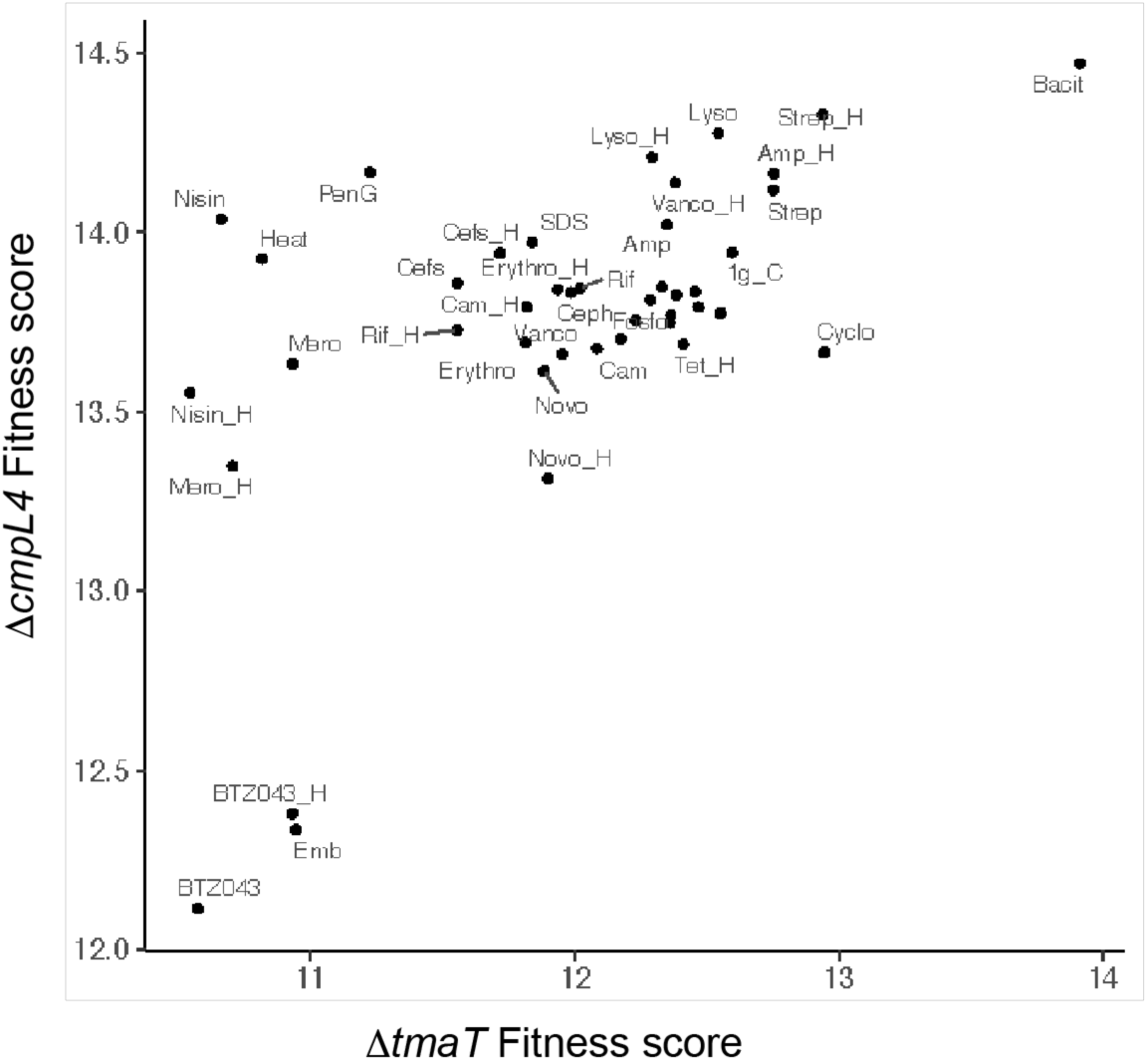
**Fitness scores correlate between *tmaT* and *cmpL4* across phenotypic profiles from Tn-seq data**^1^. Read counts were summed per gene and then normalized across all libraries with a variance stabilizing transformation from the R package DESeq2 (v1.46.0). As such, these fitness scores are dimensionless and scale with transposon insertion density. See Materials and Methods for further details.

**Fig. S5.**
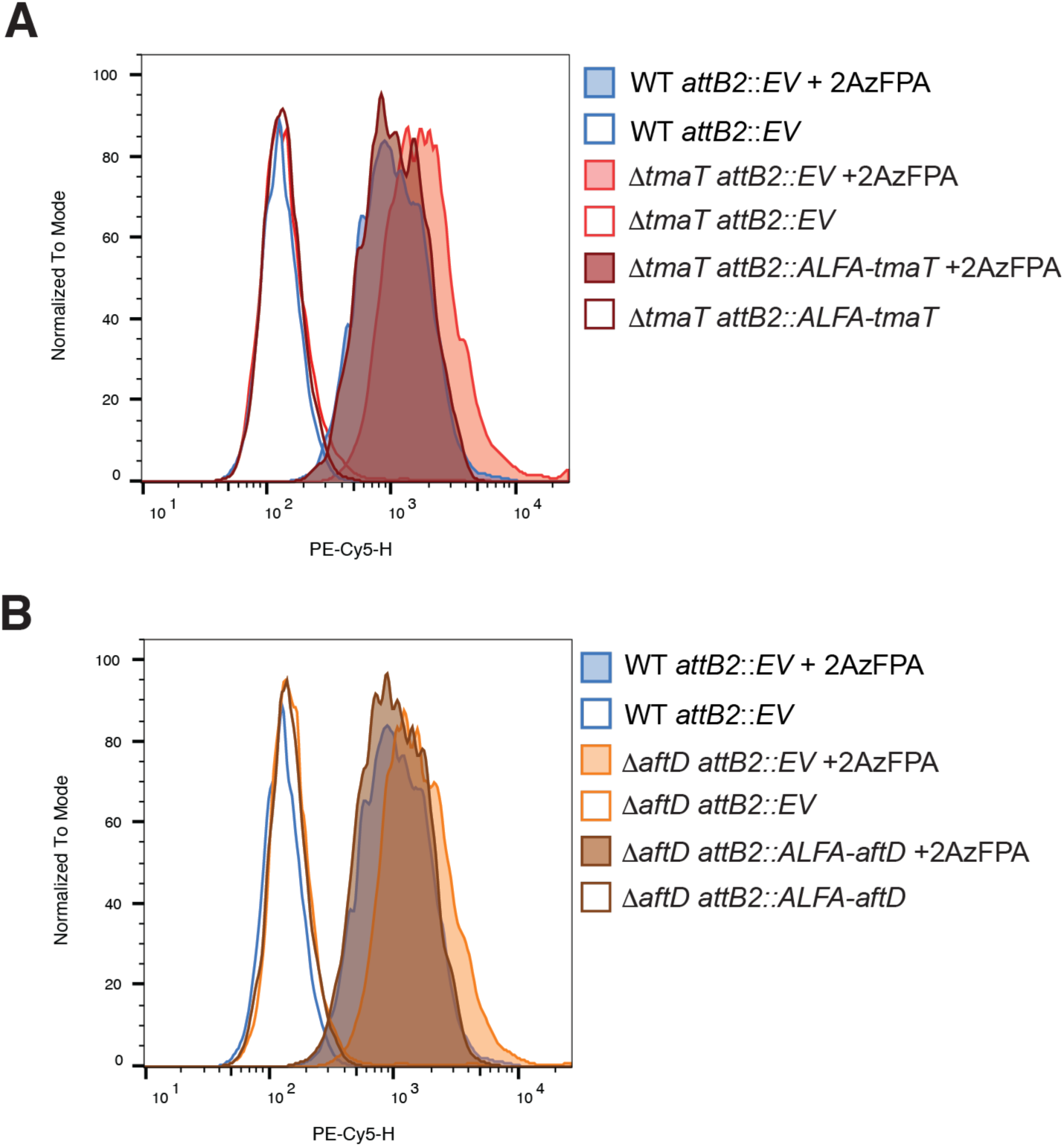
Flow cytometry histograms of 2-AzFPA incorporation in *C. glutamicum* MB001 Δ*tmaT* and Δ*aftD* strains. (A*–*B) Representative flow cytometry histograms of PE-Cy5 channel fluorescence intensity (AFDye 647) from *C. glutamicum* strains (*A*) Δ*tmaT attB2*::EV (coral), Δ*tmaT attB2*::ALFA-*tmaT* (dark red), and WT *attB2*::EV (blue). (*B*) Δ*aftD attB2*::EV (orange), Δ*aftD attB2*::ALFA-*aftD* (brown), and WT *attB2*::EV (blue) labeled with 2-Azido-(Z,Z)-farnesyl phosphoryl-β-D-arabinose (2-AzFPA). Strains were grown to OD600 0.2 in BHI, incubated with 150 µM 2-AzFPA for 4 hours at 30°C, washed, and reacted with AFDye 647 DBCO (150 µM, 1.5 h, 30°C) by copper-free click chemistry to fluorescently label incorporated 2-AzFPA. Cells were analyzed on a BD FACS Symphony A1 Analyzer; 100,000 cells were counted per sample. Filled histograms represent +2-AzFPA conditions; open histograms represent no-probe controls. Representative histograms from triplicate experiments are shown. Data were analyzed using FlowJo. See Fig. 6 for quantification of mean fluorescence intensity.

**Figure S6.**
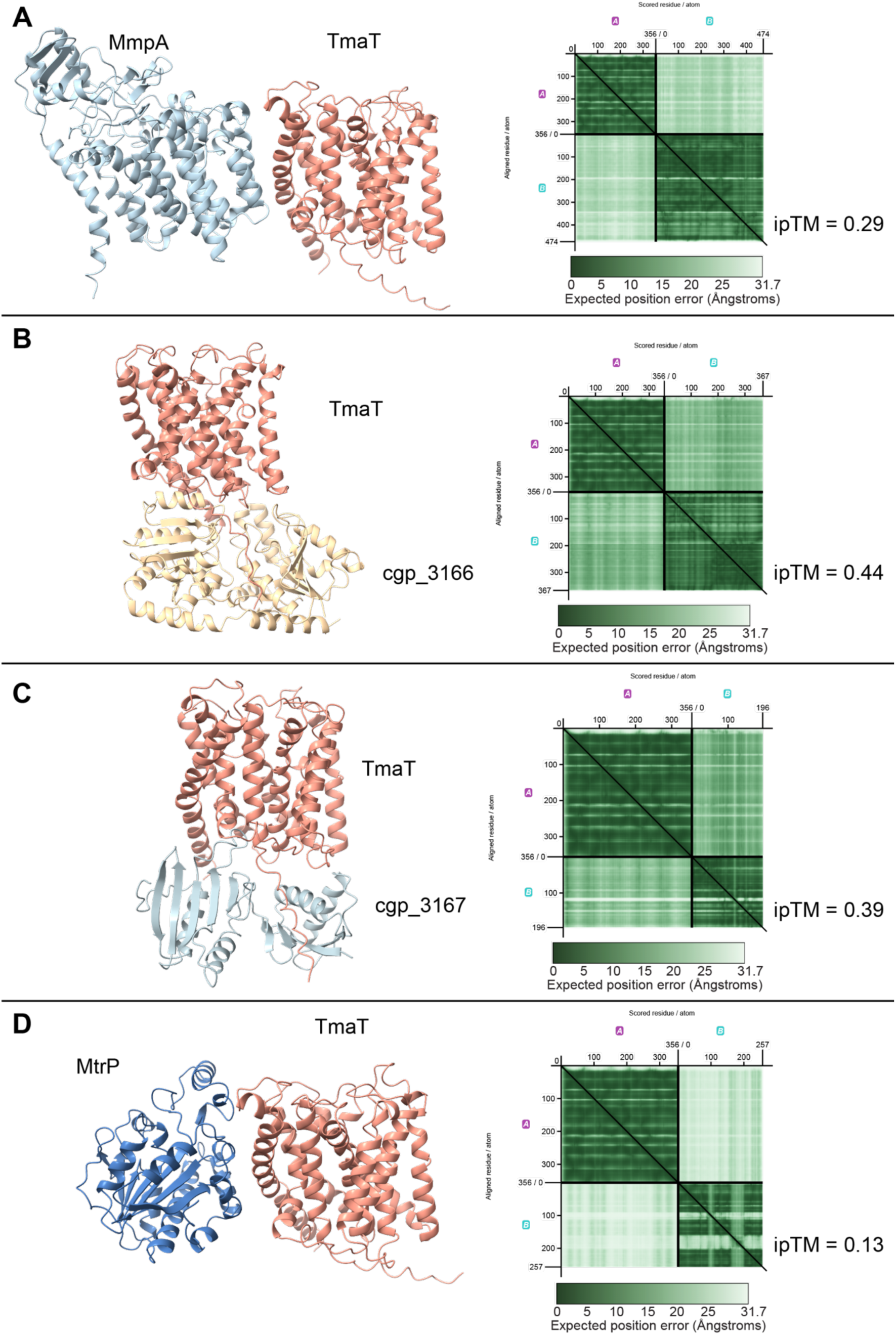
AlphaFold3 prediction of protein-protein interactions between TmaT and syntenic gene cluster proteins. (A-D) Predicted complex structures (left panels) and predicted aligned error (PAE) plots (right panels) for TmaT paired with **A)** MmpA (membrane protein), **B)** Cgp_3166, **C)** Cgp_3167, and **D)** MtrP, all from *C. glutamicum* MB001. In the PAE plots, each axis represents the residue position of each protein in the predicted complex; dark green values indicate low expected position error and high confidence in the relative orientation of those residues, while light green values indicate high positional uncertainty. Confident protein-protein interactions are reflected by dark off-diagonal blocks in the PAE plot, indicating that the predicted positions of residues in one protein are accurate relative to residues in the other protein. In all four predictions, off-diagonal blocks are uniformly light green, indicating high positional uncertainty across all inter-protein residue pairs. Interface predicted TM-score (ipTM) values, which quantify the confidence of the predicted interaction interface on a scale of 0 to 1, were 0.29 **(A)**,0.44 **(B)**, 0.39 **(C)**, and 0.13 **(D)**, all below the threshold of 0.5 required for a confident interaction prediction^2^. Taken together, these data indicate that TmaT does not form stable complexes with any of the four proteins encoded within its syntenic gene cluster under the conditions modeled. All structure predictions were generated using AlphaFold3^2^.

**Figure S7.**
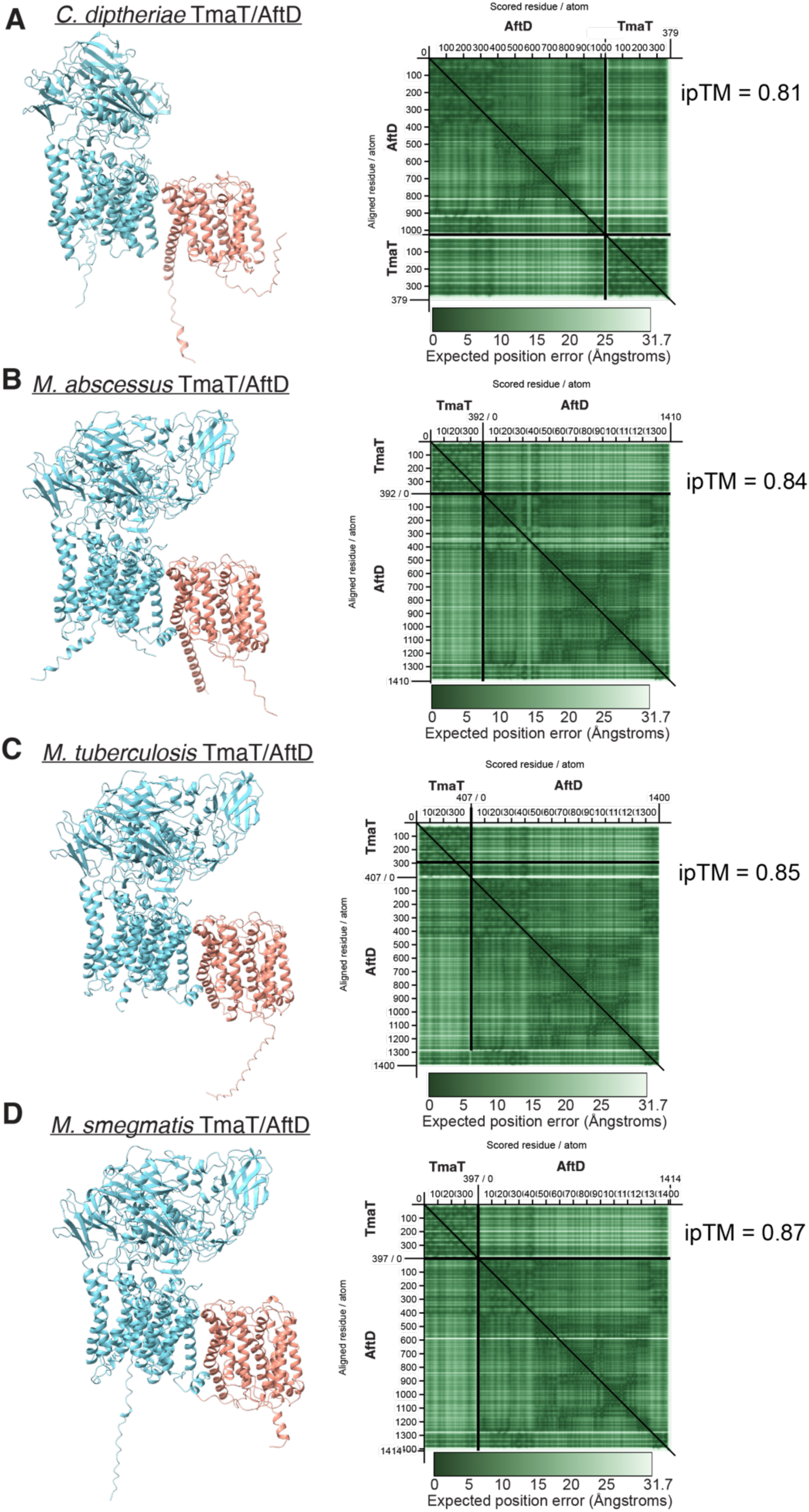
AlphaFold3 prediction of TmaT/AftD protein complex formation across Mycobacteriales. (A*–*D) Top-ranked predicted complex structures (left panels) and predicted aligned error (PAE) plots (right panels) generated using AlphaFold3 (1) and visualized using PAE-Viewer (2) for TmaT (coral) and AftD (teal) orthologs from **(A)** *Corynebacterium diphtheriae* NCTC13129 (TmaT, DIP2175; AftD, DIP2174), **(B)** *Mycobacterium abscessus* ATCC 19977 (TmaT, MAB4473c; AftD, MAB4472), **(C)** *Mycobacterium tuberculosis* H37Rv (TmaT, Rv0228; AftD, Rv0236c), and **(D)** *Mycobacterium smegmatis* mc²155 (TmaT, MSMEG_0319; AftD, MSMEG_0359). In the PAE plots, each axis represents the residue position of each protein in the predicted complex; dark green values indicate low expected position error and high confidence in the predicted relative positions of those residues, while light green values indicate high positional uncertainty. Confident protein-protein interactions are reflected by dark off-diagonal blocks in the PAE plot, indicating that the predicted positions of residues in one protein are accurate relative to residues in the other protein. Interface predicted TM-scores (ipTM) are indicated in each panel. All structure predictions were performed using AlphaFold3 and PAE plots were generated and visualized using PAE-Viewer^2,3^.

**Figure S8.**
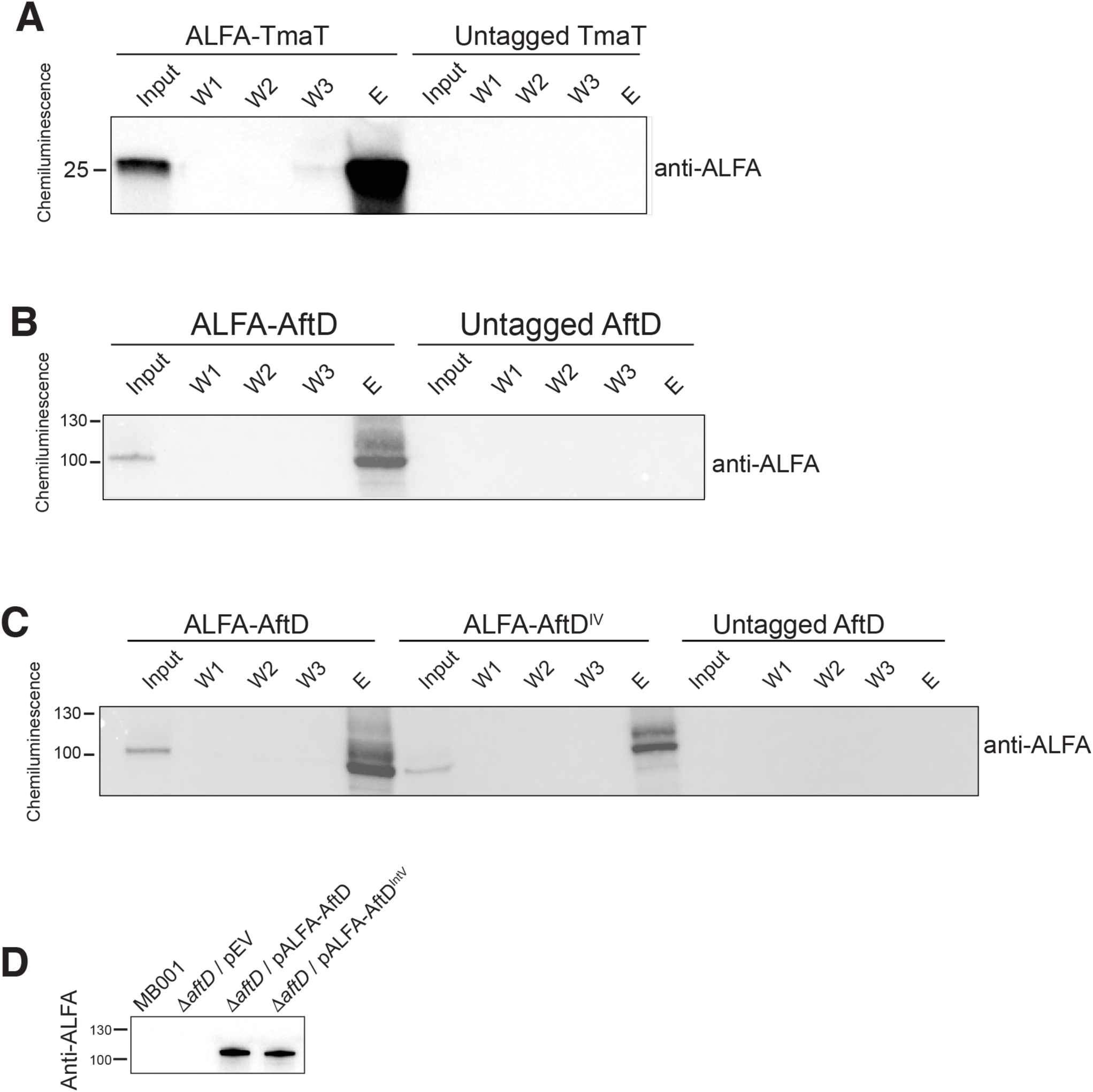
Expression controls for ALFA-tag pulldown experiments presented in. **Figs. 7 and 8. (A*–*C)** Western blots probed with anti-ALFA antibody confirming expression and ALFA-nanobody pulldown of tagged proteins. Input, wash (W1–W3), and eluate (E) fractions are shown for each condition. **(A)** ALFA-TmaT (∼25 kDa) is detected in the input and elutes following pulldown with anti-ALFA nanobody magnetic beads, confirming successful enrichment. Untagged TmaT serves as a negative control and is absent from the eluate. **(B)** ALFA-AftD (∼100 kDa) is detected in the input and elutes following pulldown. Untagged AftD serves as a negative control and is absent from the eluate. **(C)** ALFA-AftD and ALFA-AftD^IV^ (AftD interface variant) are both detected in the input and eluate fractions, confirming successful pulldown of both proteins. Untagged AftD serves as a negative control and is absent from the eluate. **(D)** Whole cell lysates from *C. glutamicum* MB001, Δ*aftD*/pEV, Δ*aftD*/pALFA-AftD, and Δ*aftD*/pALFA-AftD^IV^ probed with anti-ALFA antibody, confirming that ALFA-AftD and ALFA-AftD^IV^ are expressed at comparable levels. MB001 and Δ*aftD*/pEV lack the ALFA tag and serve as negative controls. Pulldown conditions are described in Materials and Methods.

**Fig. S9.**
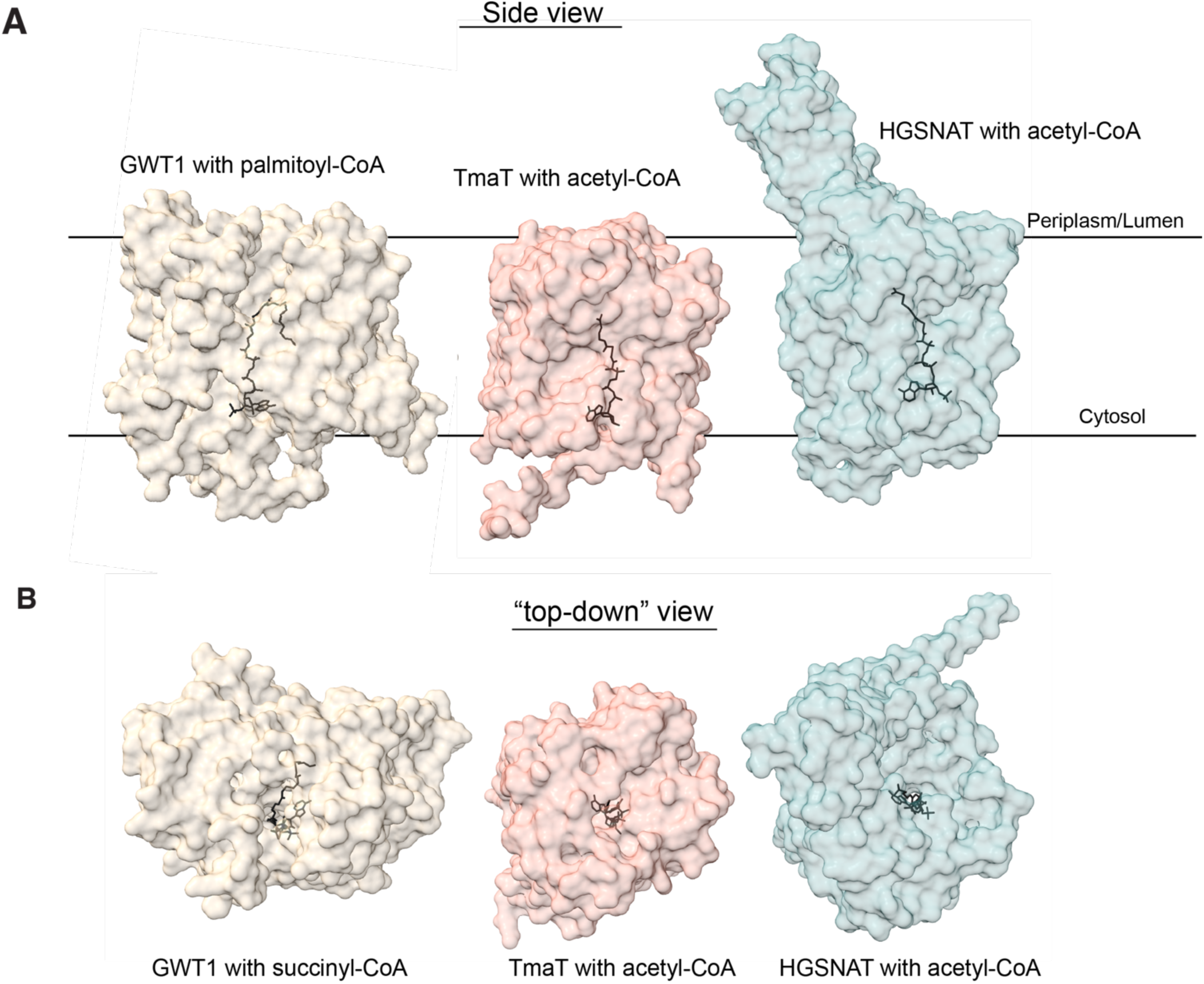
Structural comparison of TmaT with AT3 family acetyltransferases. (A*–*B) Surface representations of an AlphaFold3-predicted structure of TmaT from *C. glutamicum* MB001 co-predicted with acetyl-CoA using its SMILES representation (coral, center), compared to the solved cryo-EM structures of the AT3 family acetyltransferases GWT1 bound to palmitoyl-CoA (PDB: 8XIJ; white, left) and human HGSNAT bound to acetyl-CoA and substrate analog (PDB: 8VLG; teal, right), shown in side view **(A)** and top-down view **(B)**^4,5^. Horizontal lines in **(A)** indicate the membrane boundaries, with the periplasm/lumen above and cytosol below. Bound acyl-CoA cofactors are shown as stick representations in black. Structures were visualized using UCSF ChimeraX.

---

**Supplemental table 1:**
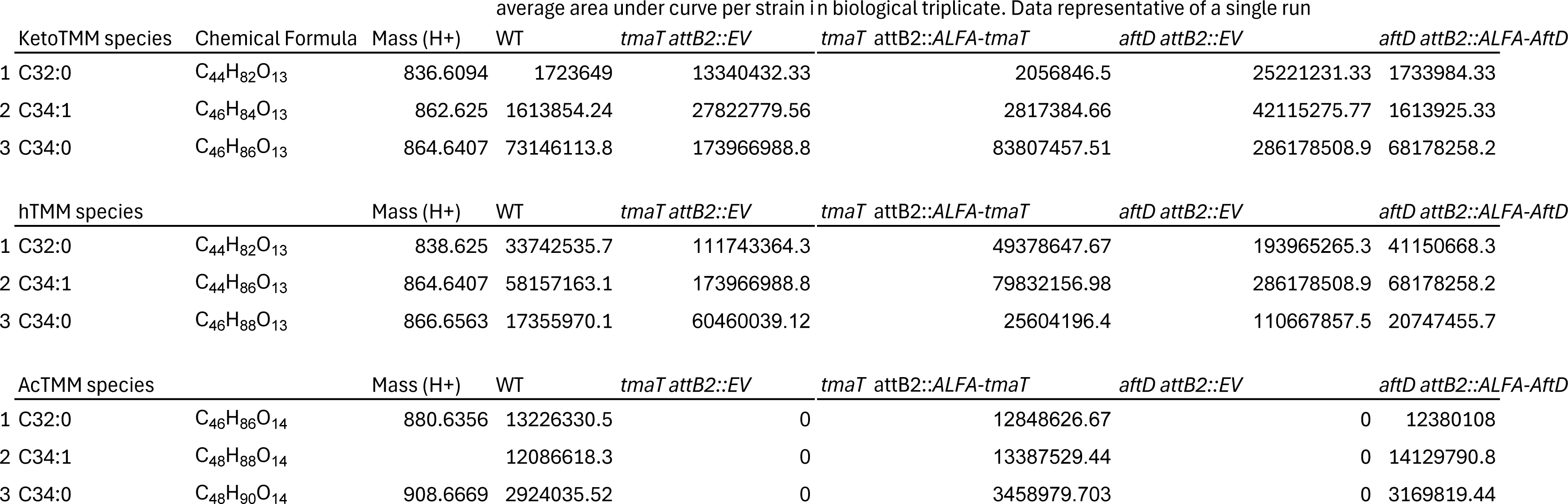
Area under curve for TMM species.

**Table S2:**
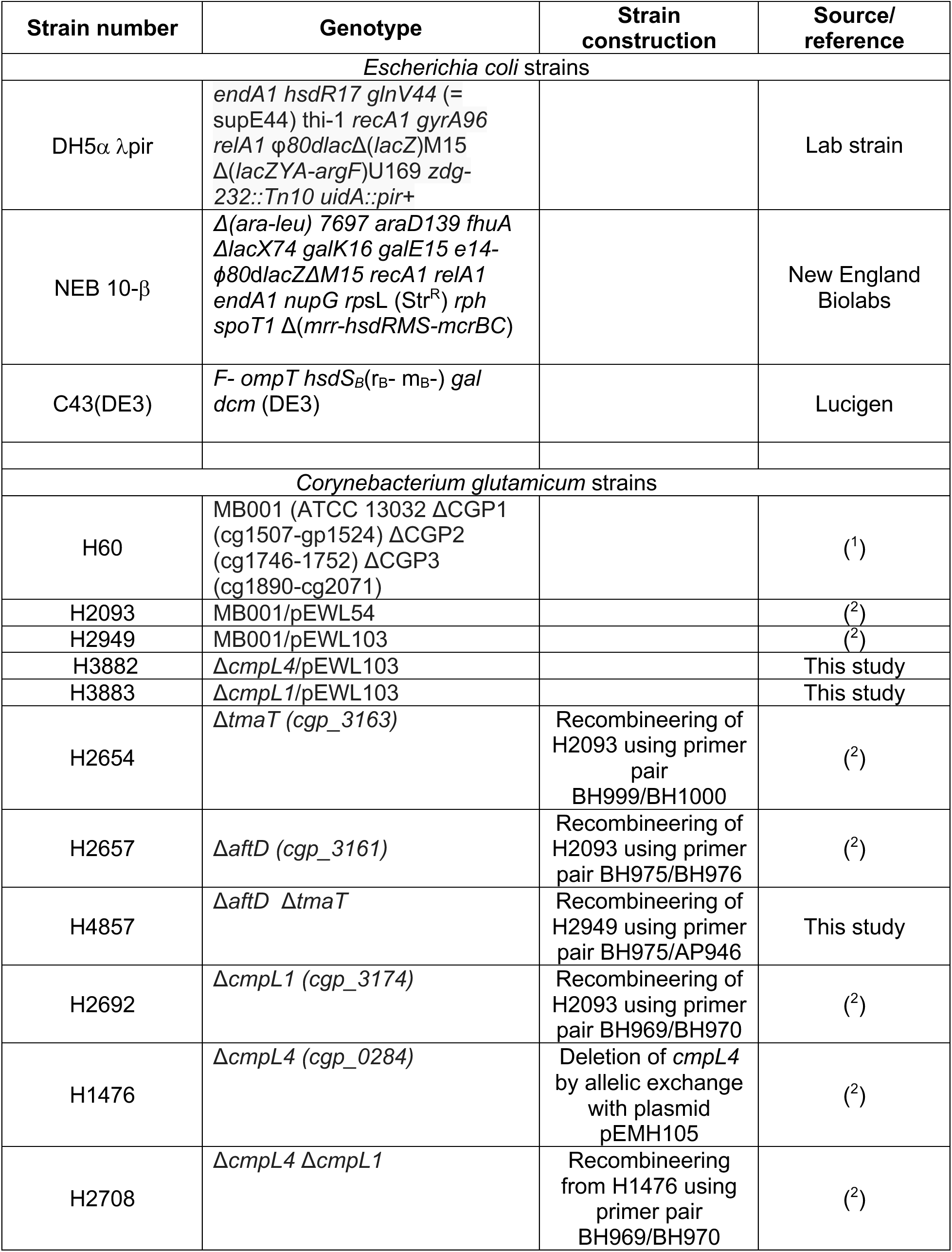
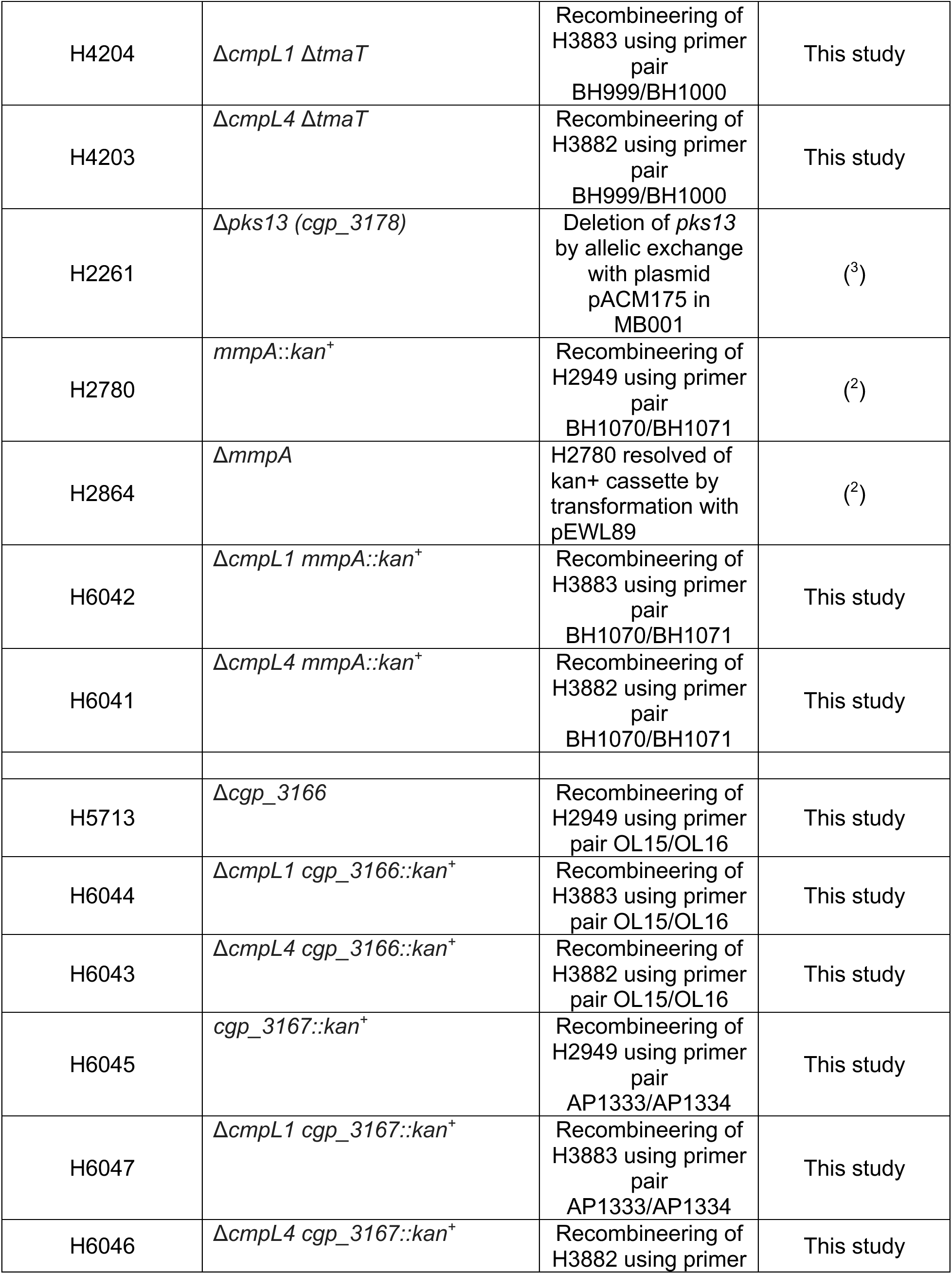
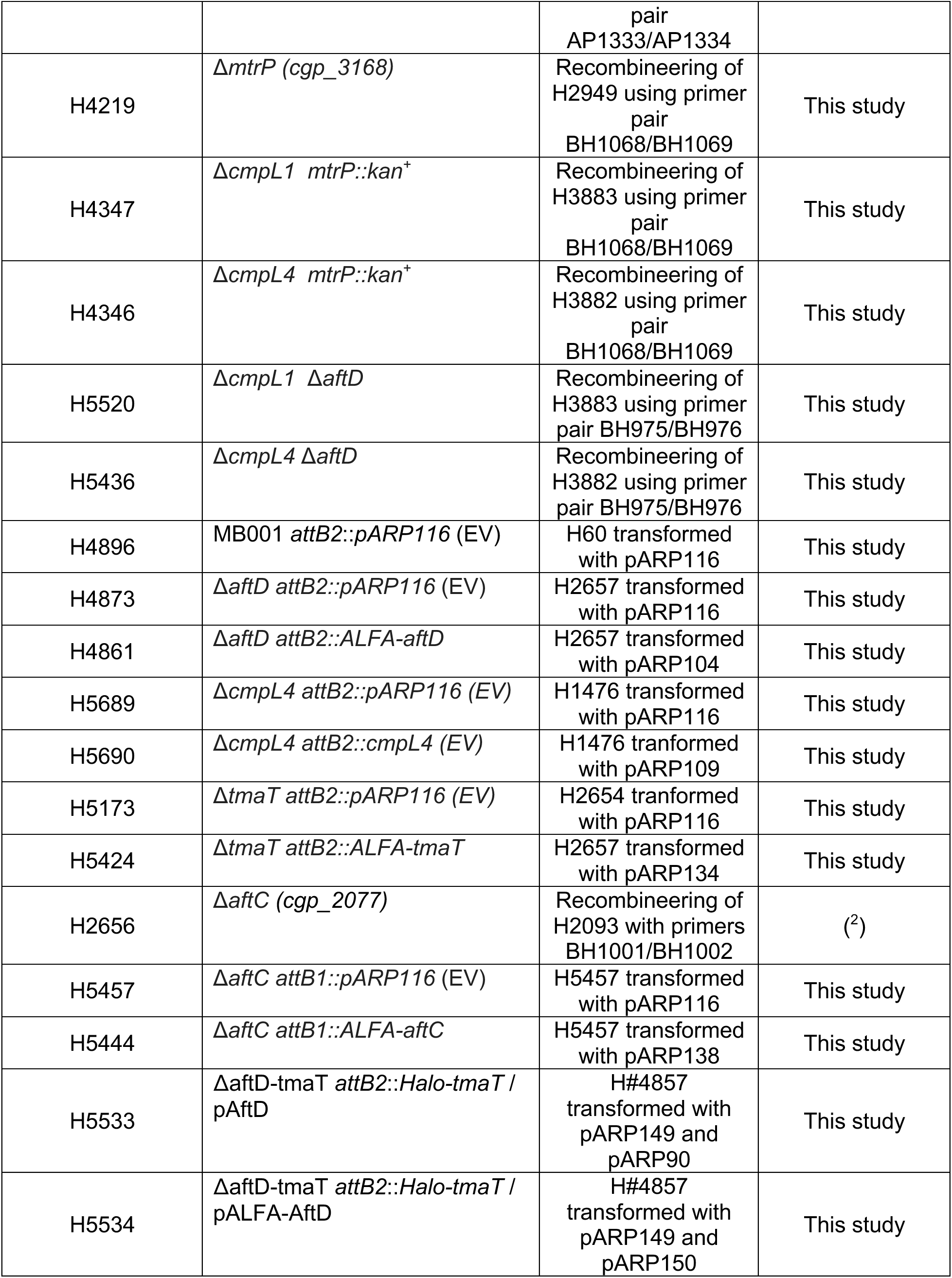
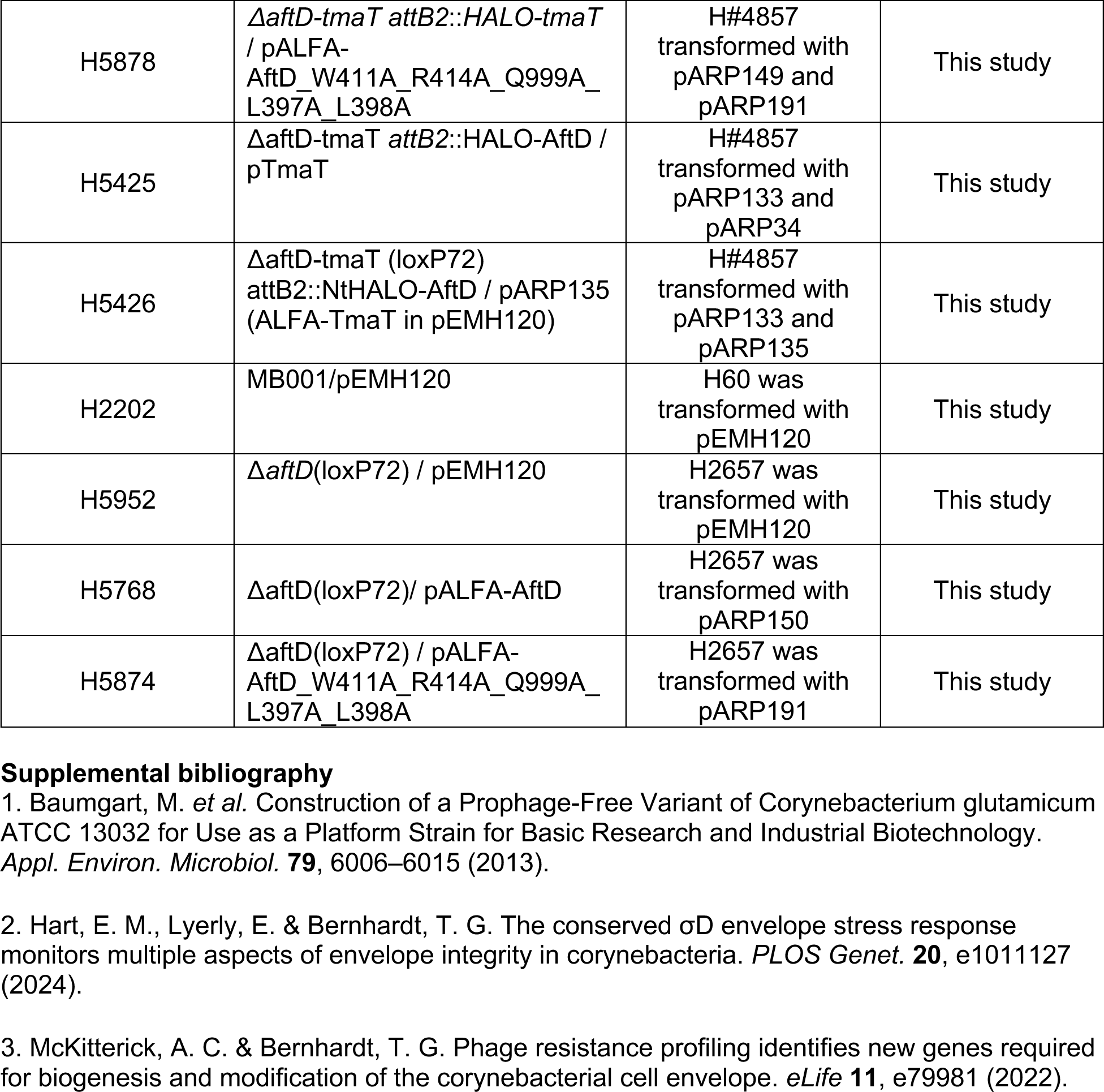
Strain list.

**Table S3:**
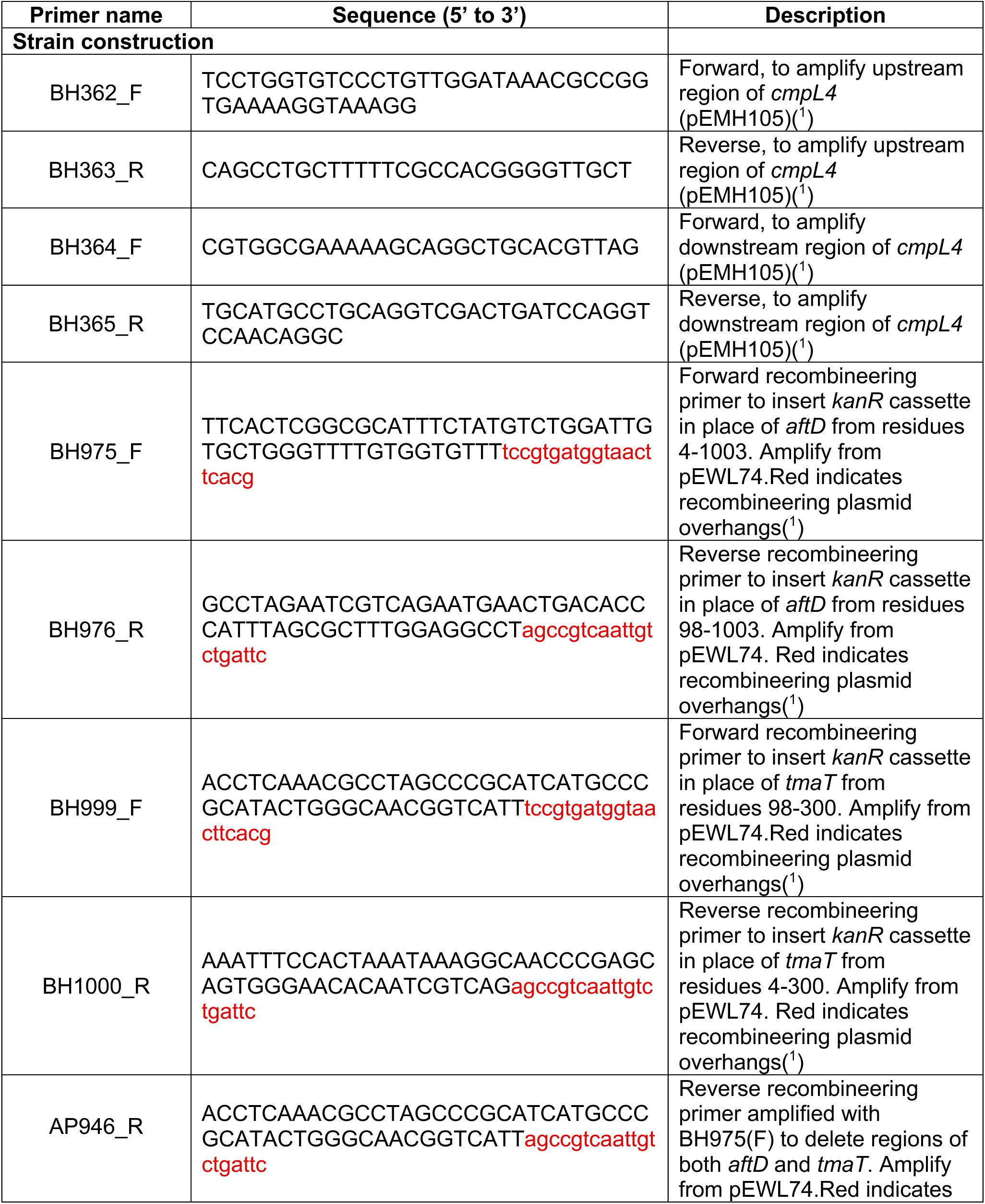
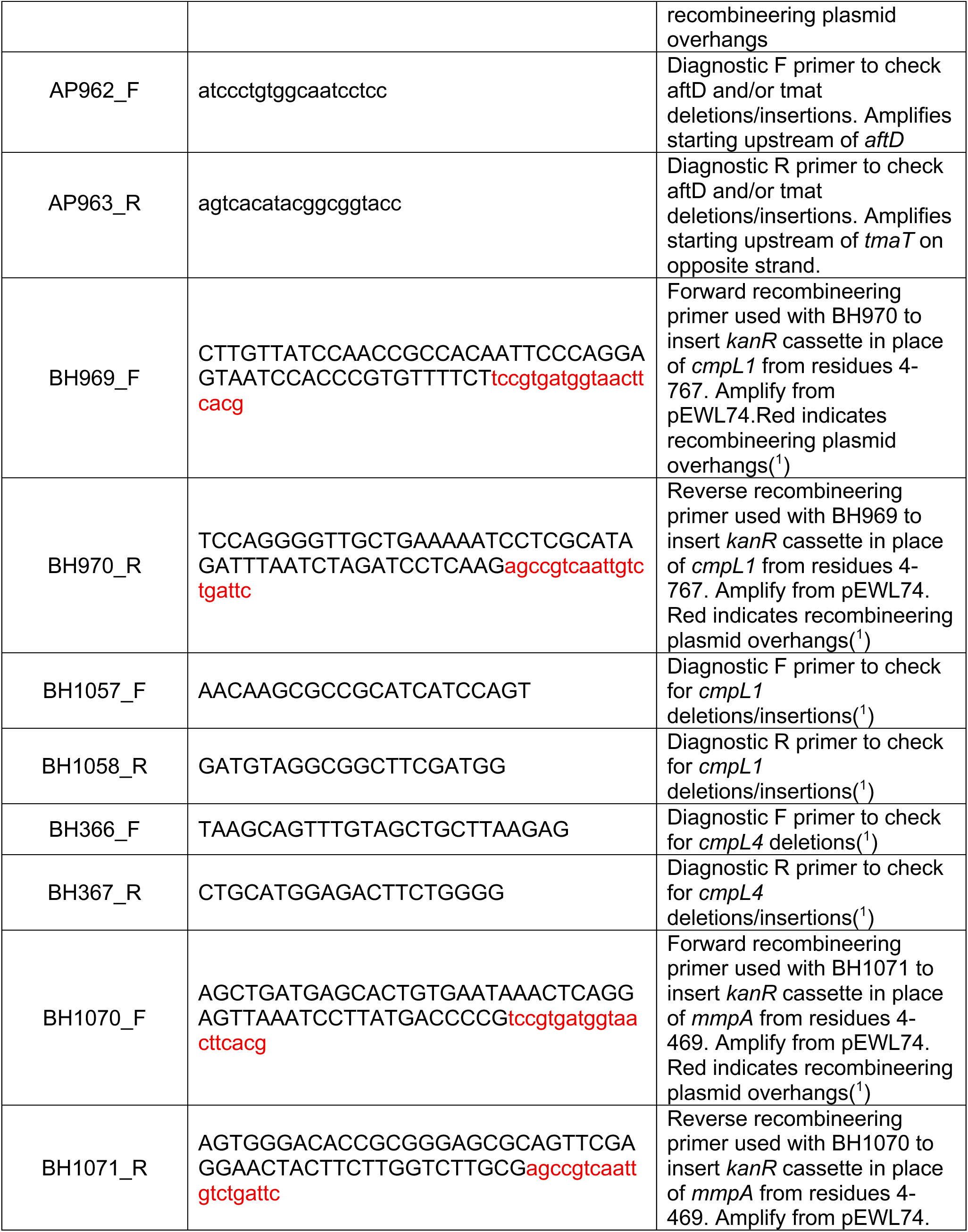
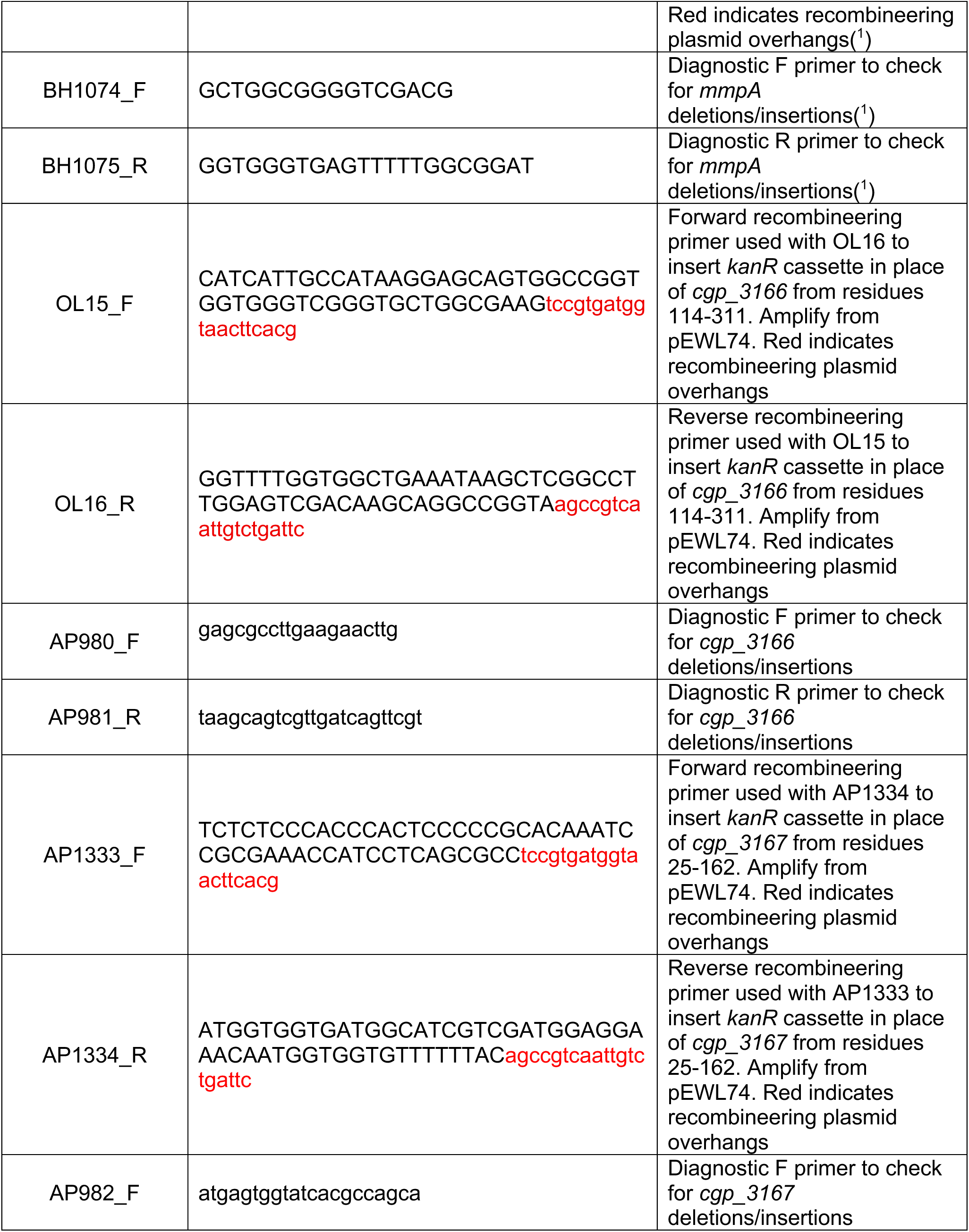
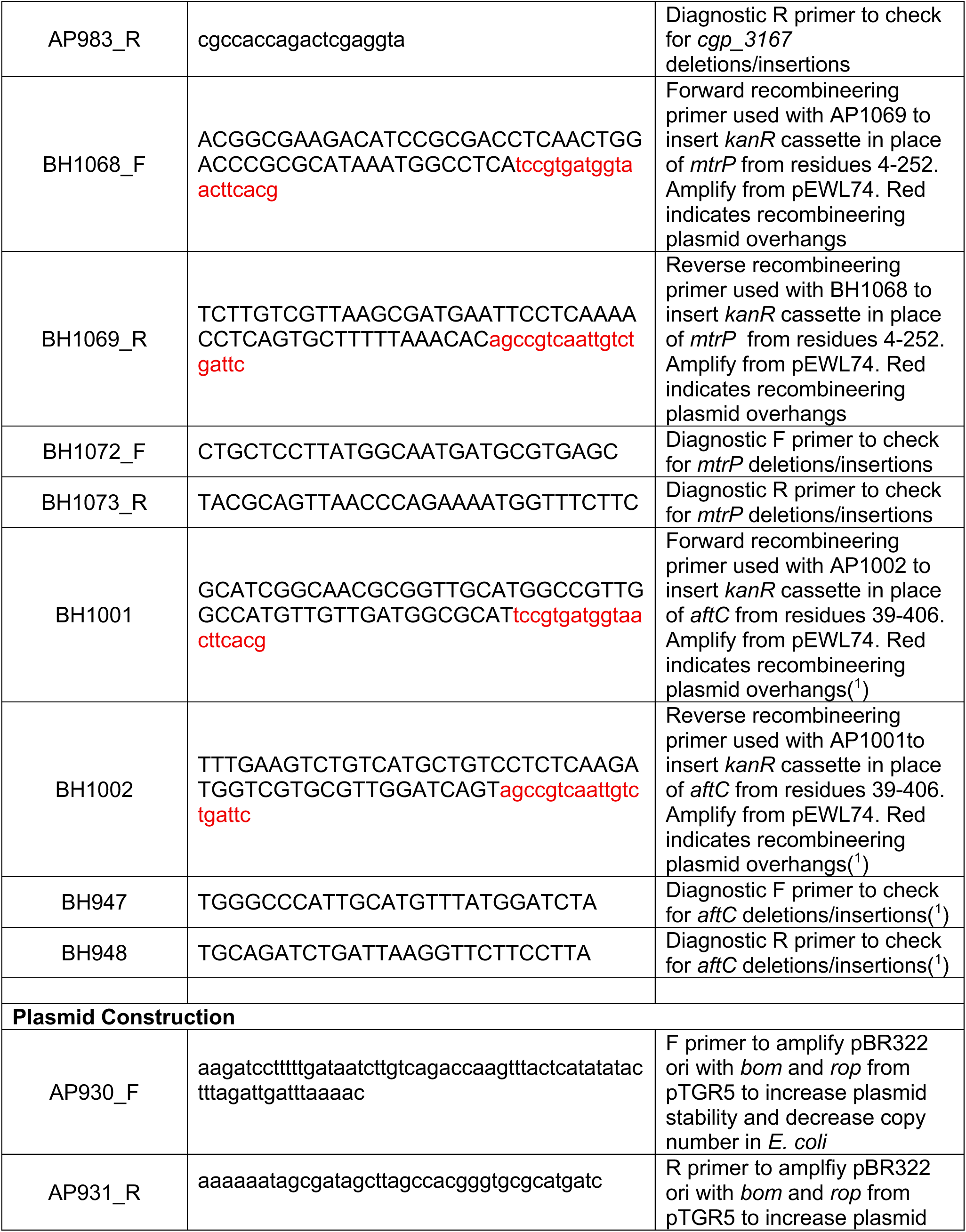
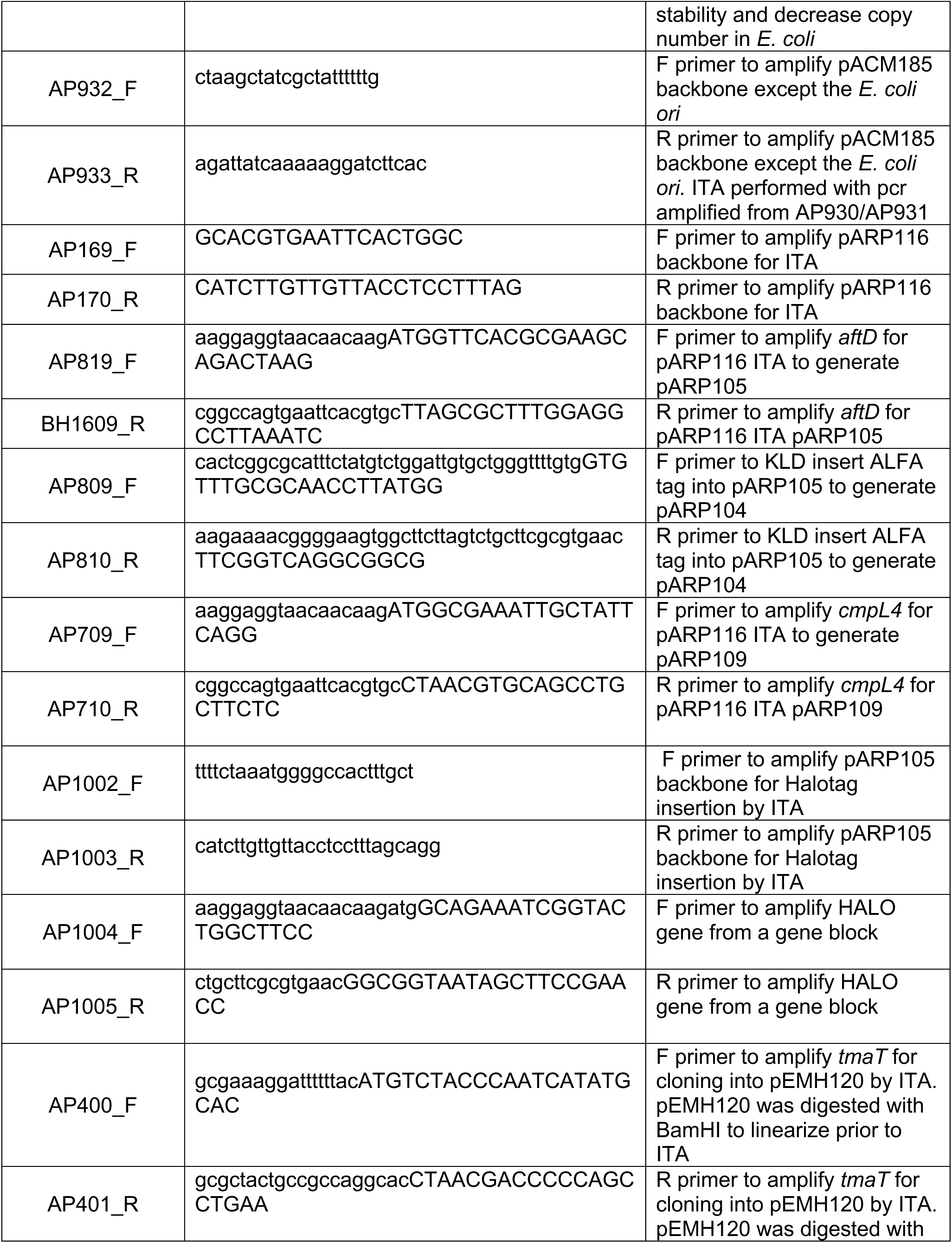
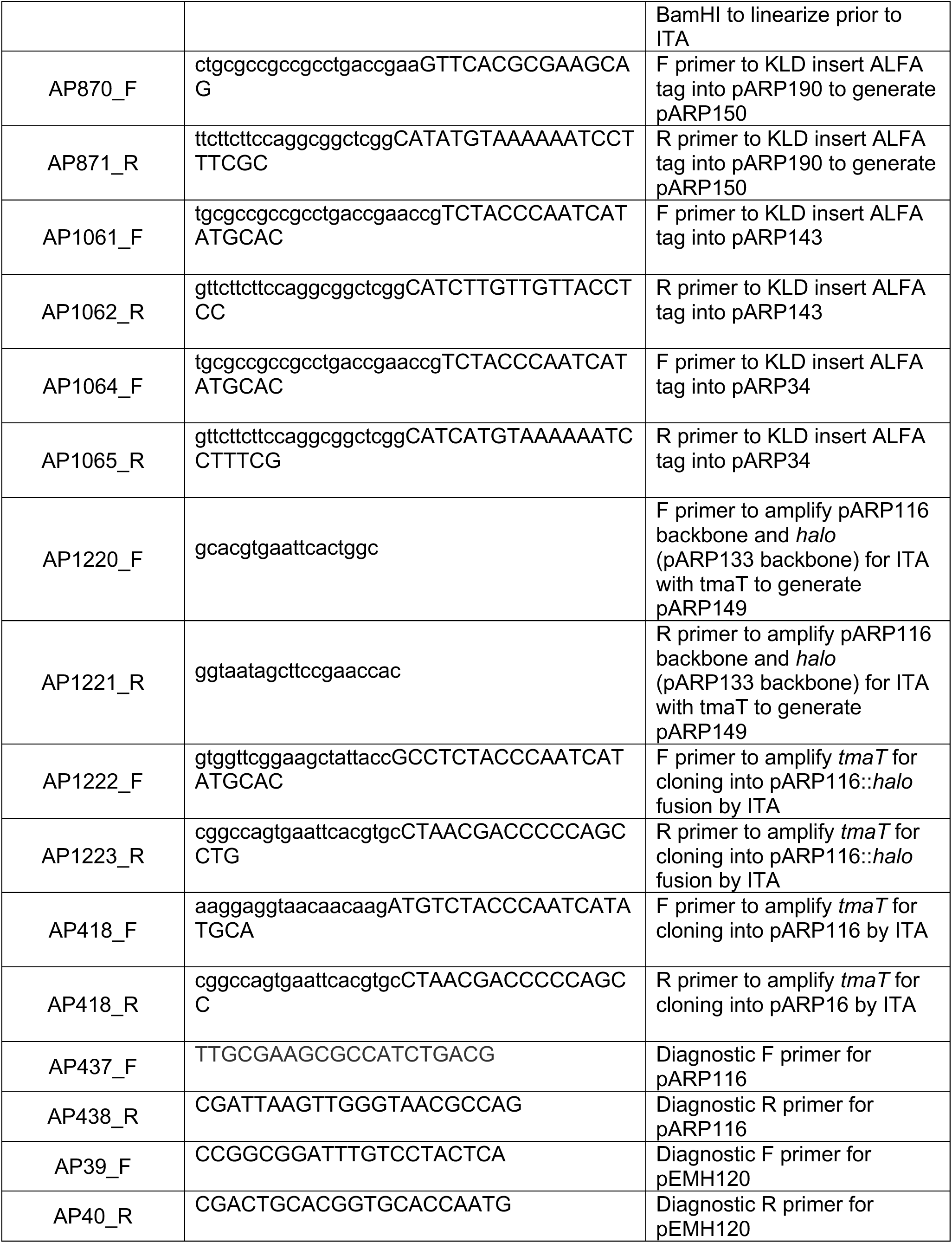
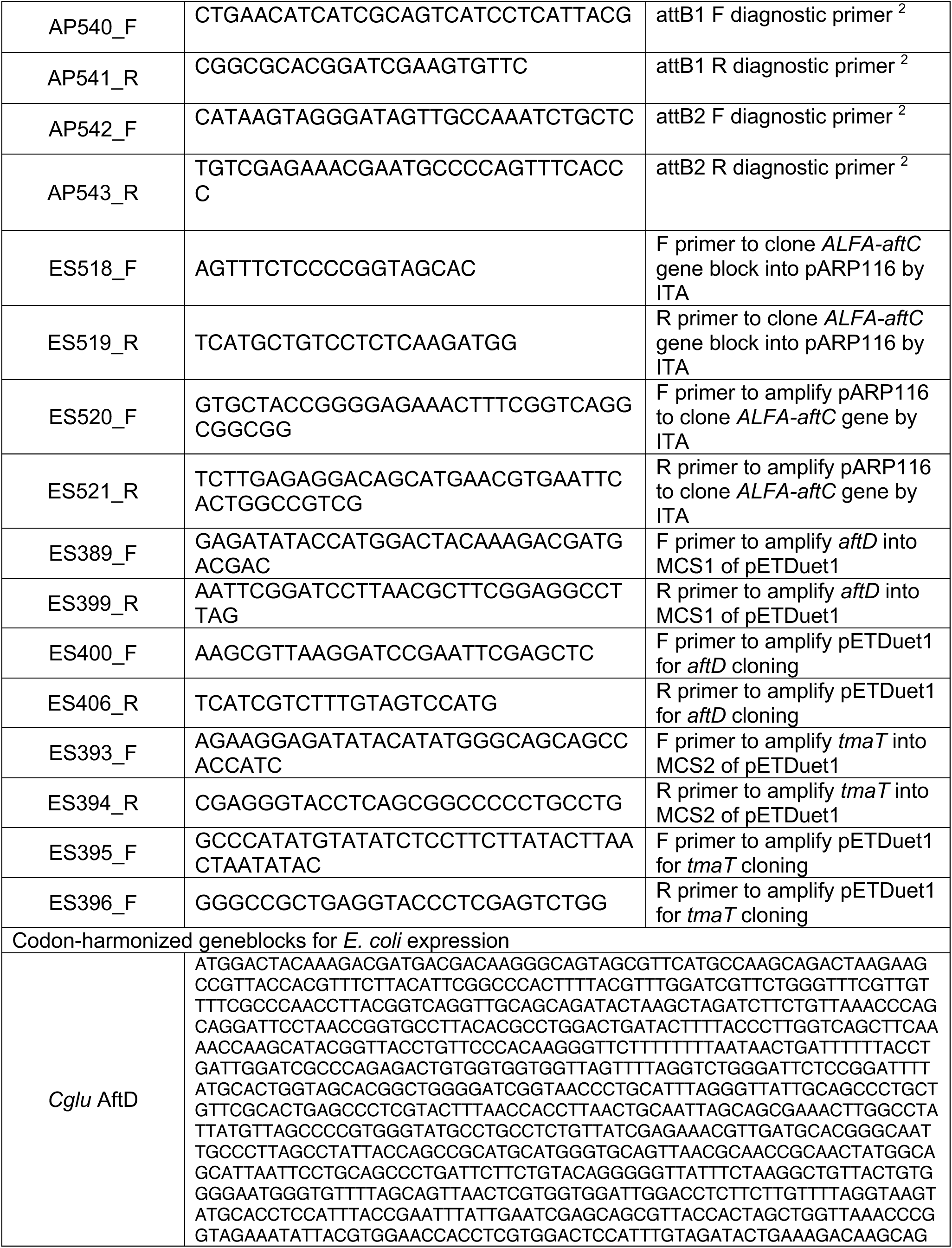
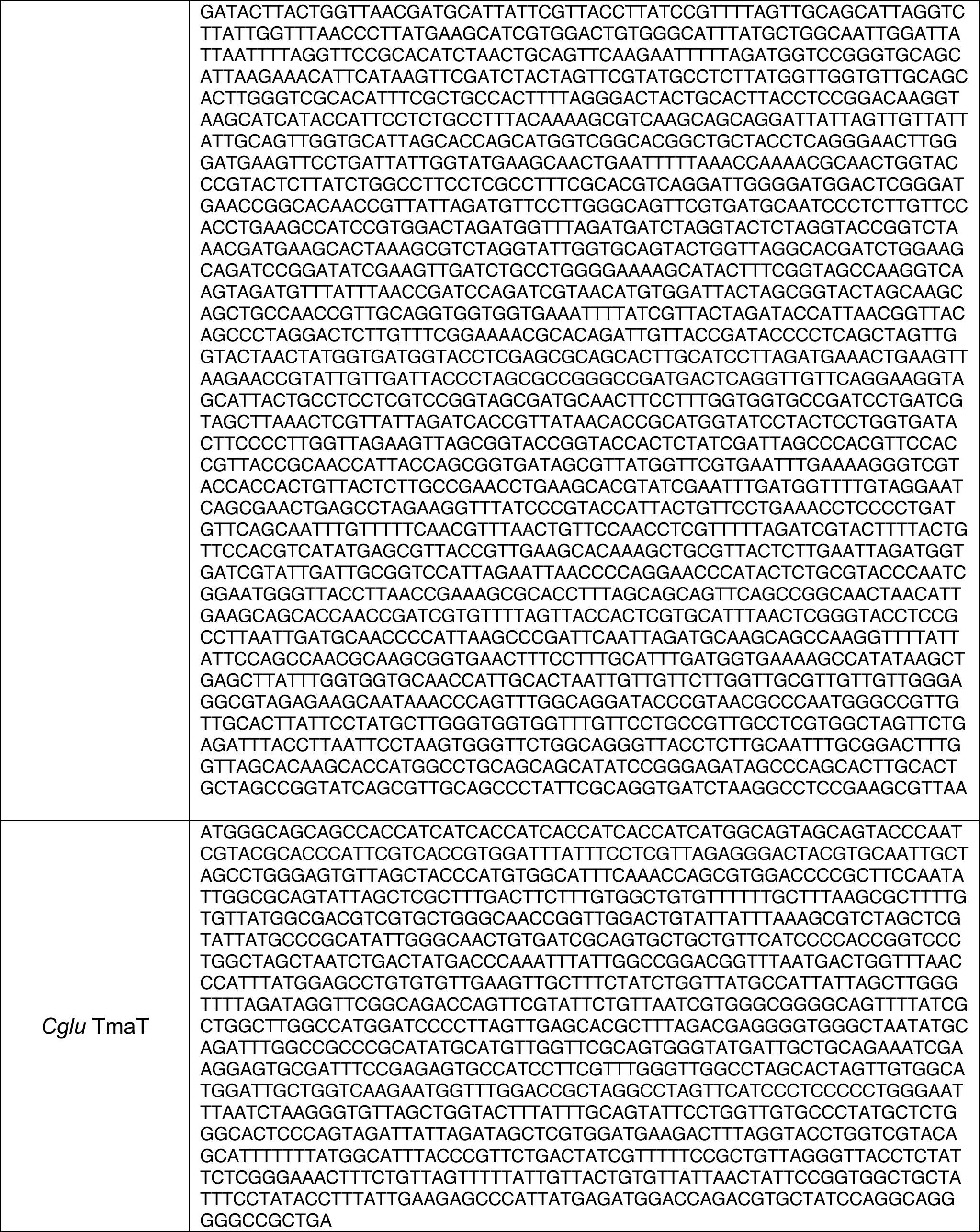
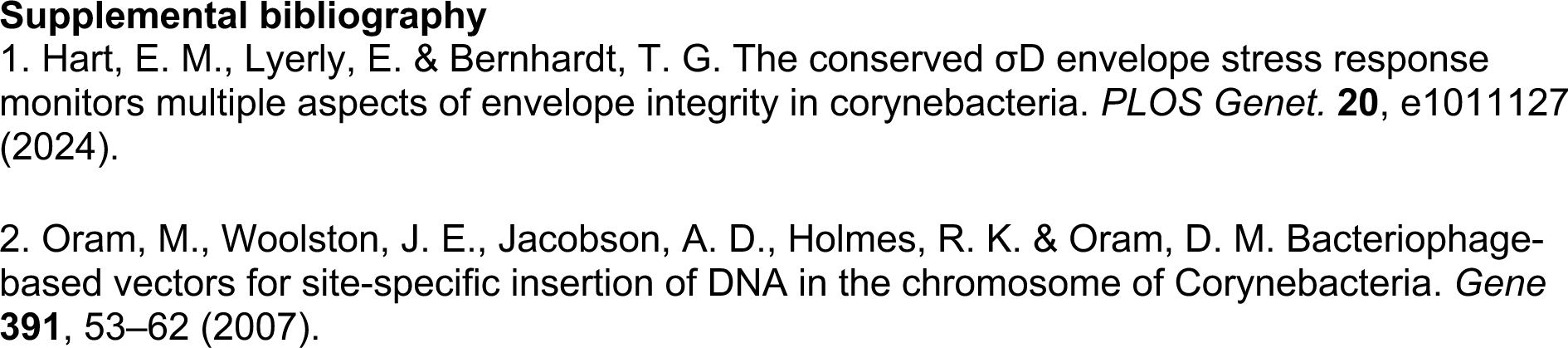
Oligonucleotides used in this study.

**Table S4:**
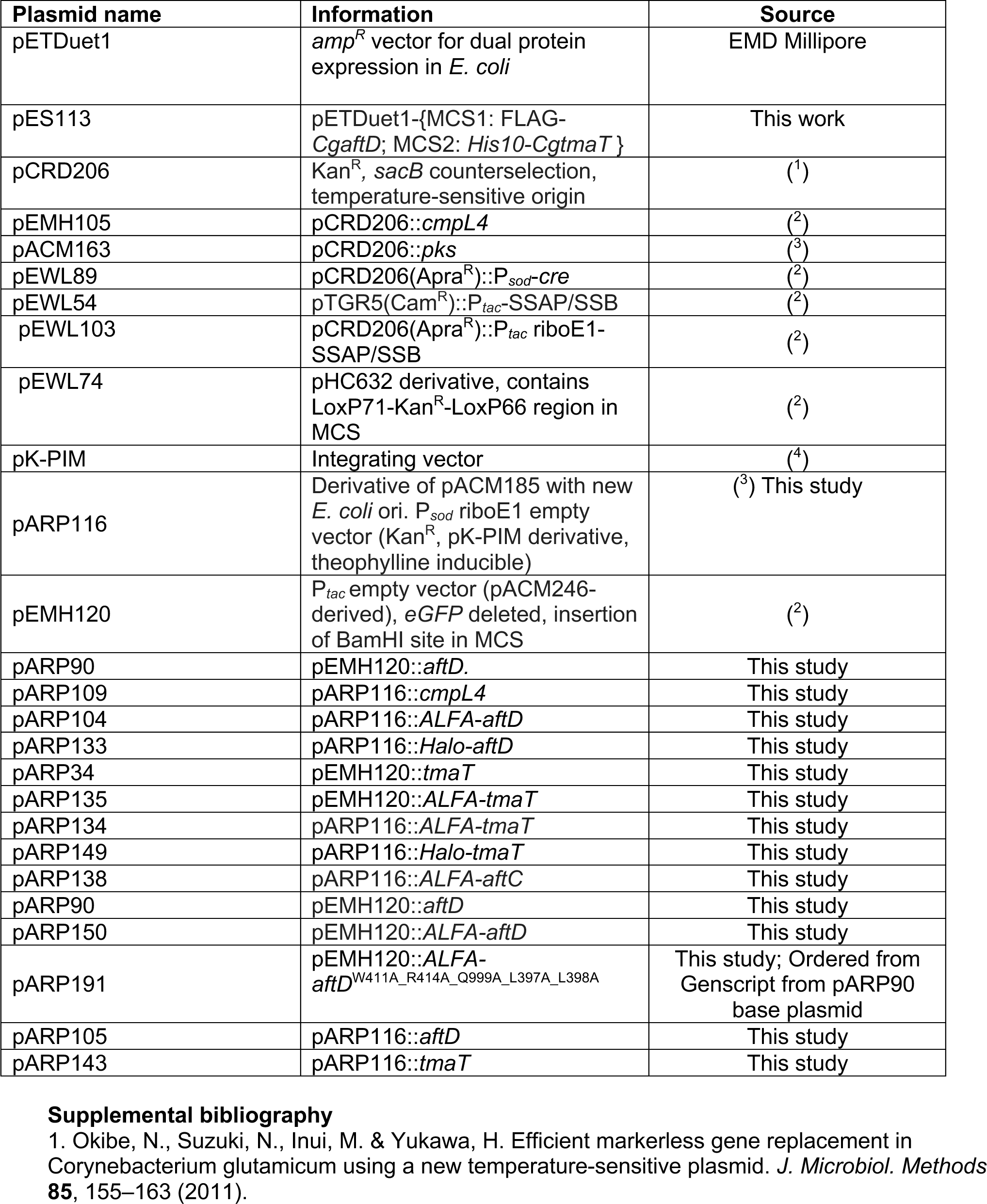
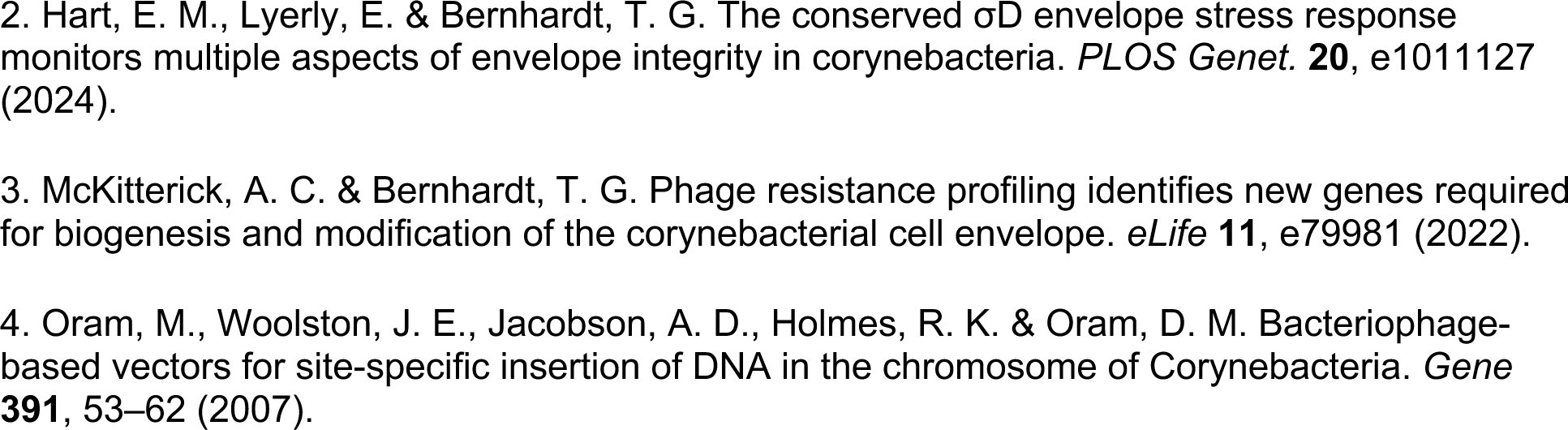
Plasmids used in this study.

## Notes

### Competing Interest Statement

The authors have declared no competing interest.

